# Comprehensive Interrogation of Synthetic Relationships in the Human DNA Damage Response

**DOI:** 10.1101/2023.08.18.553865

**Authors:** John Fielden, Sebastian M Siegner, Danielle N Gallagher, Markus S Schröder, Maria Rosaria Dello Stritto, Lena Kobel, Moritz F Schlapansky, Petr Cejka, Marco Jost, Jacob E Corn

## Abstract

The DNA damage response (DDR) is a multi-faceted network of pathways that preserves genome stability. Unraveling the complementary interplay between these pathways remains a challenge. Here, we comprehensively mapped genetic interactions for all core DDR genes using combinatorial CRISPRi screening. We discovered myriad new connections, including interactions between cancer genes and small molecule targets. We focused on two of the strongest interactions: *FEN1*/*LIG1*:*WDR48* and *FANCM*:*SMARCAL1*. First, we found that WDR48 works with USP1 to restrain overactive translesion synthesis in FEN1/LIG1-deficient cells, and that a preclinical inhibitor of USP1 specifically kills FEN1-deficient cells. Second, we found that SMARCAL1 and FANCM suppress DNA double-strand break (DSB) formation at TA-rich repeats in late replicating regions that otherwise escape into mitosis and cause nuclear fragmentation. We present fundamental insights into genome maintenance processes and our dataset provides a springboard for mechanistic investigations into connections between DDR factors and suggests multiple interactions that could be exploited in cancer therapy.

## Introduction

Numerous DNA repair pathways, collectively known as the DNA damage response (DDR), counteract diverse threats to genome integrity^1,2^. A failure to maintain genome integrity can lead to human diseases such as premature ageing and cancer^3^. Even during unperturbed cellular growth, the DDR is continually activated to deal with issues ranging from mismatches to complete DNA replication fork collapse. The roles and regulation of many individual DDR pathways have been resolved in detail^4,5^. However, the DDR is buffered by factors with overlapping functions and by pathways which can compensate for each other despite completely distinct molecular mechanisms. This can mask gene functions in essential DNA repair processes and necessitates a systematic dissection of genetic interactions in the DDR.

Mapping genetic interactions can reveal new and unanticipated biological insights^6^. Synthetic lethality, when cells tolerate the loss of individual genes but not their simultaneous removal, suggests buffering via backup pathways. Synthetic viability, when lethality caused by the loss of one gene is rescued by the simultaneous loss of another, can reveal inhibitors of rescue pathways or factors that drive cell death. Additionally, genetic interaction mapping can suggest routes to precision therapies such as when DDR-mutant cancers become hyper-dependent on other DDR pathways for their survival^7^. A prime example is the synthetic lethal interaction between *BRCA1/2*, which are frequently mutated in breast and ovarian cancers, and PARP enzymes^8,9^. It is estimated that 1/3 of all cancer types harbour mutations in DNA repair genes, raising the possibility that there are many cancer-specific vulnerabilities yet to be identified^10^.

To comprehensively map genetic interactions in the DDR, we turned to CRISPR interference (CRISPRi) dual guide screening^11^. This approach enables the simultaneous expression of two single guide RNAs (sgRNAs) to robustly silence the expression of two defined genes and has been effectively used to map genetic interactions between select pathway subsets and assign novel functions to poorly characterized genes^11^. We developed a new DDR-centric dual guide library called SPIDR (Systematic Profiling of Interactions in DNA Repair) and used it to map approximately 150,000 genetic interactions for 548 core DDR genes, revealing novel aspects of DDR biology and therapeutically relevant synthetic lethal relationships.

Mechanistic interrogation revealed two new functional relationships between WDR48 and LIG1/FEN1 and between FANCM and SMARCAL1. First, we find that cell death upon WDR48:LIG1/FEN1 co-depletion involves the deubiquitylase USP1 and is driven by impaired DNA gap ligation. This triggers unbalanced RAD18-mediated PCNA mono-ubiquitylation and inappropriate REV1-dependent translesion synthesis. Significantly, FEN1 deficiency, which is found in colorectal cancers, renders cells sensitive to a preclinical inhibitor of USP1. Second, we find that FANCM:SMARCAL1 co-deficient cells exhibit a defective response to replication stress and pronounced genome instability. This interaction does not involve the rest of the FA pathway or other enzymes that similarly to SMARCAL1 catalyse fork reversal. The translocase activities of FANCM and SMARCAL1 specifically suppress DSB formation at long TA-rich repeats in late replicating regions that might otherwise form DNA secondary structures (e.g., hairpins) leading to fork collapse.

A web-based interface to our DDR interaction network is available at http://bit.ly/41CVHP4. Altogether, our work reveals fundamental insights into how genome stability is maintained and provides a starting point for mechanistic resolution of DDR pathways and the identification of novel targets for cancer therapy.

## Results

### A dual guide CRISPRi screen maps genetic interactions in the human DDR

To systematically map genetic interactions in the DDR, we designed a combinatorial CRISPRi library targeting 548 core DDR genes, comprising all genes with the gene ontology term “DNA repair” (GO:0006281). We performed the screen in the absence to exogenous DNA damage to query interactions between pathways required for normal cell function. We opted for a CRISPRi rather than CRISPR nuclease-based screen to avoid generating confounding DSBs, a source of DNA damage that could activate the DDR. By not inducing complete loss-of-function of target genes, we reasoned that CRISPRi would also enable us to identify genetic interactions involving essential DNA repair genes and to better model DDR deficiencies found in cancer, which frequently arise from hypomorphic mutations or epigenetic silencing resulting in reduced expression^12,13^.

The sgRNAs targeting each gene were selected from the human CRISPRi-v2 library^14^. The final library consisted of at least 2 sgRNAs targeting each gene, each paired with every other sgRNA (Fig. 1a). For each essential gene, we included a mismatched variant of the top sgRNA that was empirically validated or predicted to confer partial knockdown^15^. Each targeting sgRNA was also paired with 15 non-targeting sgRNAs, and 225 non-targeting-only dual sgRNAs pairs were included as negative controls. In total, the library consisted of 697,233 dual sgRNA constructs targeting 149,878 gene pairs (Supplementary Table 1). Each sgRNA combination was uniquely synthesized on an oligonucleotide chip and cloned into a dual-sgRNA lentiviral expression vector^16^. For ease of reference, we called this library SPIDR (Systematic Profiling of Interactions in DNA Repair).

**Figure 1.**
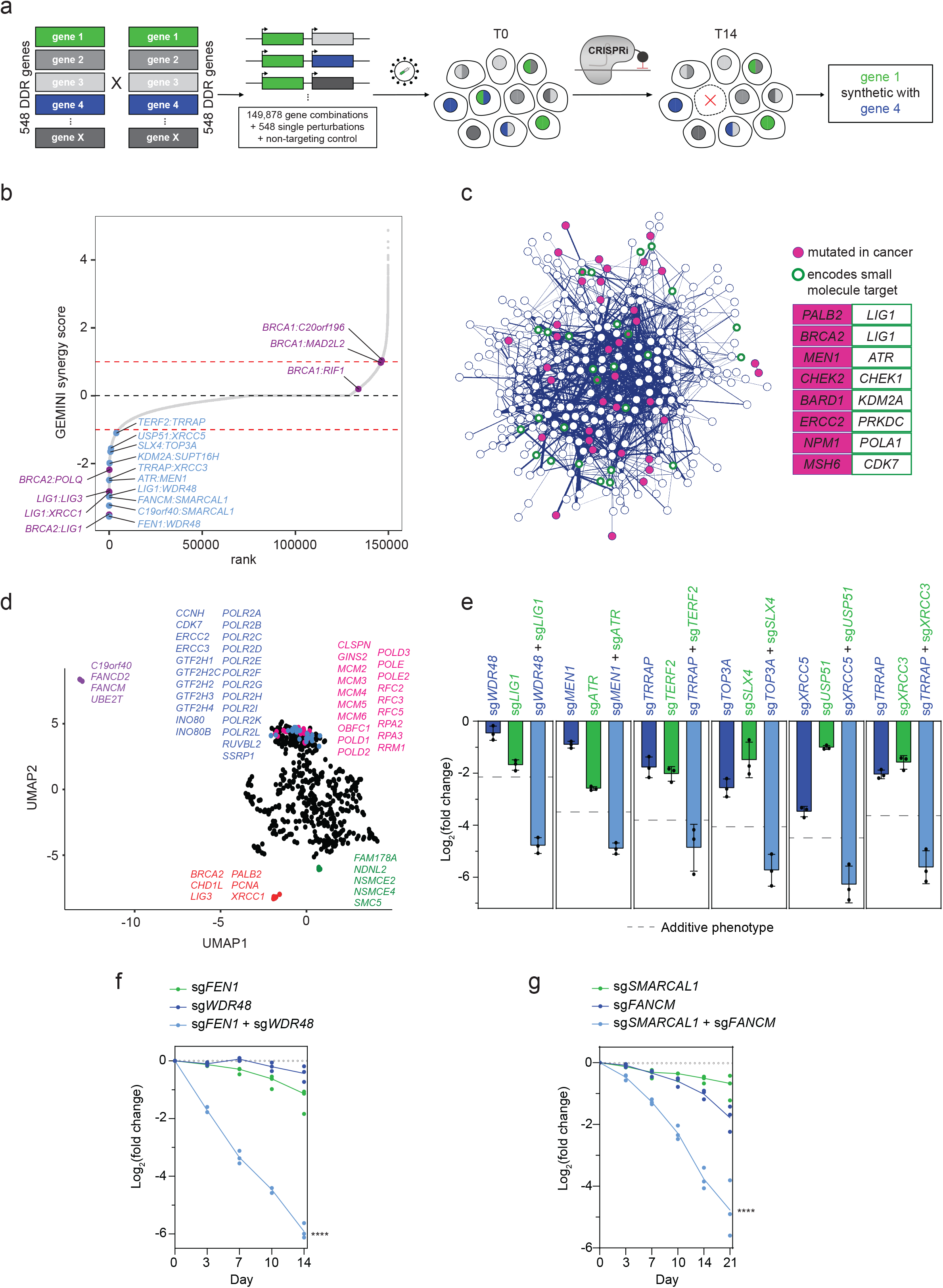
Comprehensive combinatorial screening identifies synthetic relationships between DDR factors: **a,** Guide RNAs targeting 548 core DDR genes (GO:0006281) are systematically combined into a dual-guide lentiviral library termed SPIDR (Systematic Profiling of Interactions in DNA Repair). Guide RNA samples were taken at time zero (T0) and day 14 (T14). The screen was performed with two technical duplicates. **b,** Rank-ordering all pairwise interactions by sensitive GEMINI score identifies known synthetic relationships (magenta) and nominates new candidates (light blue). Red dashed lines indicate a sensitive GEMINI score of +/-1. **c,** Synthetic relationships filtered at GEMINI score ≤ -1.0 are represented as a complex network, with the strength of the interaction determining the width of the edge. Nodes are colored based on whether they are mutated in cancer (magenta filled, COSMIC tier1 and tier2 genes), small molecule targets (green rings, DGIDb targets), or both (magenta filled and green rings). Select genetic interactions that may be therapeutically actionable (one member mutated in cancer, the other targeted by an existing small molecule) are highlighted. **d,** UMAP projecting the full set of interactions of each DDR gene with all other DDR genes. Several DDR gene neighborhoods with known, shared functions are highlighted. The UMAP was generated using the strong GEMINI score. **e,** Experimental validation of 6 novel synthetic lethal interactions identified by the SPIDR screen. Experiments were performed in triplicates and the percentage of each cell population was quantified by FACS and normalized to the corresponding untransduced cells. The log_2_ fold changes on day 14 relative to day 0 are shown. The additive phenotype (dashed line) was calculated by summing the individual depletion phenotypes. Complete survival time courses are shown in Extended Data Fig. 2c-i. **f,** Dual-color flow cytometry survival time course of the *FEN1*:*WDR48* interaction. **g,** Dual-color flow cytometry survival time course of the *FANCM*:*SMARCAL1* interaction. For f and g, 3 technical replicates from one experiment are shown and the experiments were repeated at least twice. *P*-values were calculated using two-way analysis of variance (ANOVA) between cells expressing both sgRNAs and each sgRNA alone (\*\*\*\**p* ≤ 0.0001).

Using the SPIDR library, we performed a growth-based screen in RPE-1 *TP53*-knockout (KO) cells, which have an otherwise intact DDR genetic background. We generated a clonal cell line stably expressing catalytically inactive Cas9 fused to a Krüppel associated box (KRAB) transcriptional repressor domain^17^. Cells were transduced with the lentiviral library and timepoint 0 (T0) cells were harvested 96 hours post-transduction. The final timepoint was harvested 14 days later (T14) (Fig. 1a). Next-generation sequencing and sgRNA quantification identified sgRNA pairs whose knockdown improves or inhibits cell proliferation (Extended Data Fig. 1a-b).

After confirming a strong correlation between screen replicates, we used both replicates to calculate growth phenotypes (Extended Data Fig. 1c). To benchmark the phenotypes of single gene perturbations, we assessed combinations in which the same perfectly-matched (i.e. not mismatched) sgRNA was in the A and B position. The abundance of all non-targeting:non-targeting combinations was closely distributed around zero at T14 relative to T0, indicating they did not cause major effects on cell proliferation (Extended Data Fig. 1d). Known essential genes were depleted, including components of the minichromosome maintenance (MCM) and the general transcription factor (TF) IIH complexes. sgRNAs targeting negative growth regulators, such as SAMHD1 and TAOK1, were enriched at T14 (Extended Data Fig. 1d). The screen therefore captures phenotypes of individual gene knockdowns, enabling us to infer the differences in the observed effects of gene pairs relative to that expected from the individual gene knockdowns.

To identify synergistic interactions that exceed additive single gene effects, we used a variational Bayesian pipeline (GEMINI) that is specialized for the discovery of genetic interactions from CRISPR screening data^18^. The differences between the observed effect of sgRNA pairs and their inferred additive phenotype are used to calculate a GEMINI score that indicates the strength of a synthetic lethal or synthetic viable interaction (Supplementary Table 2).

The SPIDR DDR screen identified many previously reported genetic interactions (Fig. 1b). For example, repression of *LIG1* or *POLQ* was synthetic lethal with loss of HR factors, such as *BRCA2*. Cells with impaired HR rely on gap ligation by LIG1 and alternative DSB repair pathways, such as POLQ-mediated MMEJ, for survival^19,20^. *LIG1* also had genetic interactions with both *LIG3* and *XRCC1,* which compensate for LIG1 in Okazaki fragment ligation^21,22^. We also detected many previously unknown genetic interactions, some of which had stronger GEMINI scores than known ones (Fig. 1b).

By calculating genetic interactions using the mismatched sgRNAs for empirically essential genes (defined as single knockdown causing a growth phenotype of LFC less than -3), we identified an expanded set of interactions that included synthetic lethal relationships between otherwise essential factors. These included previously reported interactions between *BRCA2* and *RAD9A* as well as *CHEK1* and *TOP3A* (Extended Data Fig. 1e-f). Depletion of FA pathway components (e.g., FANCD2 and FANCM) was synthetic lethal with repression of *RPA1*, consistent with a known activation of the FA pathway in response to RPA1 depletion^23^. We observed a genetic interaction between *NSMCE1* and *NSMCE4A*, two members of the SMC5/6 complex, as well as between *SHFM1* and *XRCC2*, both of which have roles in HR (Extended Data Fig. 1f). Thus, the mismatched sgRNAs in the SPIDR library uncovered synthetic lethal interactions that were otherwise obscured by growth arrest or cell death upon strong depletion of essential genes.

We also re-discovered known synthetic viable genetic interactions between *BRCA1* and *TP53BP1*, *MAD2L2*/*REV7*, and *C20orf196*/*SHLD1* (Fig. 1b, Extended Data Fig. 1g). BRCA1 antagonises the NHEJ-promoting protein 53BP1 to promote HR-mediated DSB repair in S phase and loss of the 53BP1-REV7-SHLD axis restores HR in BRCA1-deficient cells^24–29^.

To explore whether our screen identified genetic interactions that could be therapeutically exploited, we mapped tier 1 and 2 cancer genes from COSMIC and small molecule targets from the Drug Gene Interaction Database onto our DDR network^30,31^. This identified synthetic lethal interactions which connect cancer genes with small molecule targets (Fig. 1c). For example, *ERCC2* mutations, found in approximately 20% of muscle-invasive bladder cancers, might render these cancers sensitive to DNA-PKcs inhibition^32^. Thus, our DDR network suggests possible avenues for therapeutic intervention and future drug development, one of which we further explored in subsequent mechanistic experiments.

We asked whether our comprehensive interrogation of DDR interactions revealed distinct gene clusters with similar functions based on similarities in their interaction profiles. We used uniform manifold approximation and projection (UMAP) to embed the interaction of each DDR gene with all other DDR genes as signatures in a lower-dimensional space. Genes involved in similar aspects of the DDR tightly grouped together (Fig. 1d). For example, Fanconi Anemia (FA) genes involved in interstrand crosslink (ICL) repair and components of the SMC5/6 complex formed distinct clusters.

We individually tested several of the strongest novel genetic interactions using an orthogonal flow cytometry-based method (Extended Data Fig. 2a-b)^11^. An sgRNA targeting each member of a synthetic lethal pair was separately cloned into a lentiviral vector that either co-expresses BFP or GFP and packaged into lentivirus. RPE-1 *TP53* KO dCas9-KRAB cells were co-transduced at the same multiplicity of infection (MOI) with both sgRNA-containing viruses, allowing us to monitor the proliferation of co-depleted cells relative to singly depleted cells. We validated interactions between *LIG1*:*WDR48*, *ATR*:*MEN1, TERF2*:*TRRAP, SLX4*:*TOP3A, XRCC5*:*USP51, XRCC3*:*TRRAP, SUPT16H*:*KDM2A, FEN1*:*WDR48*, and,

*FANCM*:*SMARCAL1* (Fig. 1e-g, Extended Data Fig. 2b-i). In all cases, the co-depleted cells exhibited a profound and synergistic proliferation defect relative to the corresponding single depletions. These genetic interactions span a wide range of potential molecular functions, from DSB repair (*XRCC5*:*USP51*) and recombination intermediate processing (*SLX4*:*TOP3A*) to epigenome maintenance (*SUPT16H*:*KDM2A*). *TRRAP*, a member of the Tip60 histone acetyltransferase complex that is recruited to DSBs, has synthetic lethal relationships with the telomere shelterin complex component, *TERF2*, and the RAD51 paralog and prostate cancer risk gene *XRCC3*^33,34^.

For mechanistic interrogation, we prioritised synthetic lethal gene pairs with 1) the strongest GEMINI scores, 2) one factor that exhibits strong interactions with multiple members of the same pathway and/or 3) one gene that is frequently mutated in cancer. Filtering our dataset using these criteria led us to two unanticipated modules: *WDR48* with *LIG1* and *FEN1,* both Okazaki fragment repair factors, and *SMARCAL1* with two genes that encode physically interacting proteins, *FANCM* and *C19orf40*/*FAAP24* (Fig. 1f, g, Extended Data Fig. 2j).

### WDR48-USP1 regulate repair pathway choice at post-replicative gaps

During single-strand (ss)DNA break repair and Okazaki fragment ligation, the FEN1 endonuclease cleaves 5’ DNA flaps and LIG1 seals the resulting ssDNA gap^35,36^. Cells lacking FEN1 or LIG1 accumulate DNA gaps, nicks, and/or flaps and rely on both HR and XRCC1-LIG3 for survival^19,21,22^. Our dataset recapitulated known genetic interactions between *LIG1* and HR (*PALB2* and *BRCA2*) and backup Okazaki fragment ligation factors (*LIG3* and *XRCC1*). In addition, we observed striking novel synthetic lethal relationships between both *LIG1* and *FEN1* with *WDR48*, which had GEMINI scores stronger or equivalent to *LIG1*:*LIG3* and *LIG1*:*PALB2* (Fig. 1b, e, f, Extended Data Fig. 2j, 3a).

WDR48 stimulates the activity of the deubiquitylases USP1, USP12, and USP46 and we individually tested whether they were involved in *WDR48’s* strong genetic interaction with *FEN1*^37^ (Extended Data Fig. 3b). Whereas knockdown of either USP12 or USP46 had no effect on the viability of FEN1-depleted cells, USP1 depletion was synthetic lethal with loss of FEN1. This suggests that the aberrant accumulation of a ubiquitylated substrate of WDR48-USP1 might underlie the *FEN1*/*LIG1*:*WDR48* genetic interactions. The WDR48-USP1 complex has multiple important substrates that regulate genome stability, including FANCD2-FANCI during crosslink repair and PCNA during translesion synthesis^38,39^

We hypothesized that DNA gaps that accumulate in the absence of LIG1 or FEN1 might stimulate PCNA mono-ubiquitylation by the RAD18 E3 ubiquitin ligase to facilitate the recruitment of specialised TLS polymerases. Without WDR48-mediated regulation, the low processivity and high-error rate of these polymerases could lead to slow DNA replication and genome instability^40^. Consistent with this, treatment with a FEN1 inhibitor (FEN1-IN-1; FEN1i) led to increased PCNA mono-ubiquitylation and DSB signaling in WDR48-depleted cells (Extended Data Fig. 3c). We used three orthogonal approaches to test the hypothesis that ssDNA gaps trigger unrestrained PCNA mono-ubiquitylation, hyperactive TLS, and cell death in FEN1/LIG1:WDR48 co-depleted cells.

First, we performed competitive growth assays to assess the sensitivity of WDR48, RAD18, and co-depleted cells to FEN1i. Co-depletion of RAD18 completely reversed the high sensitivity of WDR48-depleted cells to FEN1i (Extended Data Fig. 3d). We then generated triple-knockdown cells that express sgRNAs targeting *LIG1* and *WDR48,* plus either a non-targeting sgRNA or one targeting *RAD18*. Strikingly, RAD18 depletion completely rescued *LIG1*:*WDR48* synthetic lethality (Fig. 2a). RAD18 co-depletion also conferred a competitive growth advantage to LIG1-deficient cells in the presence of the selective WDR48-USP1 inhibitor ML-323 (Extended Data Fig. 3e)^41^.

**Figure 2.**
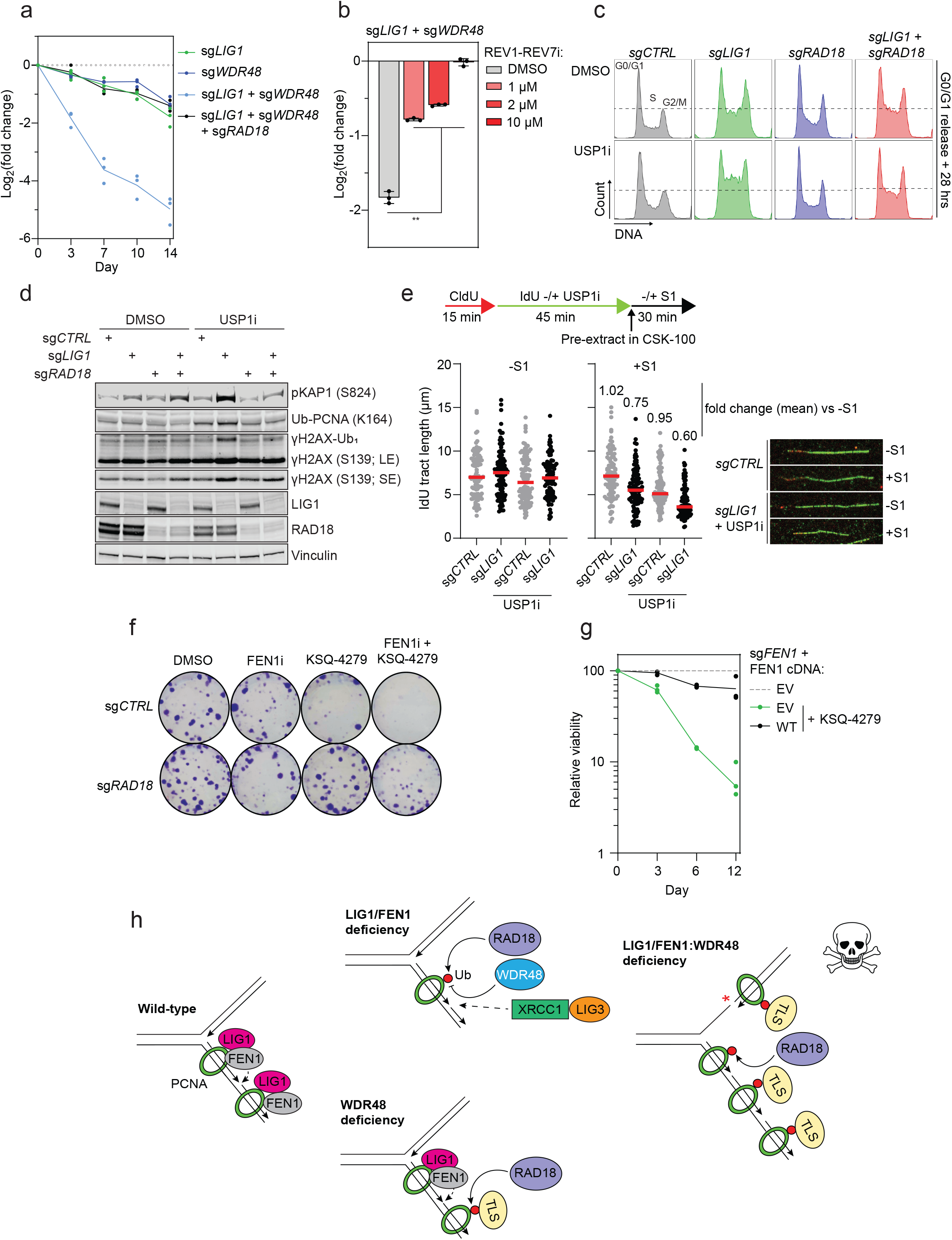
WDR48-USP1 counteracts RAD18 to promote S phase progression and suppress gap formation: **a,** Dual-color flow cytometry survival time course of the *LIG1:WDR48* interaction in RPE1 *TP53* KO dCas9-KRAB cells expressing a non-targeting or *RAD18*-targeting sgRNA. A representative experiment performed in technical triplicates is shown and was repeated once. **b,** Dual-color flow cytometry assay of LIG1:WDR48-depleted cells that were treated with DMSO or the indicated doses of the REV1-REV7 interface inhibitor, JH-RE-06 (REV1-REV7i) for 72 hours (hrs). Error bars represent mean ± s.d. \*\**p* ≤ 0.01; paired Student’s *t* test. **c,** Representative flow cytometry profiles of DNA content (DAPI) analysis in cells transduced with the indicated sgRNAs 28 hrs after release from G0/G1. G0/G1 arrested cells were released into the cell cycle in the presence of DMSO or the USP1 inhibitor ML-323 (USPi; 30 μM). Quantification of Fig. 2c and all timepoints are shown in Extended Data Fig. 3f, g. **d,** Western blot of cells transduced with the indicated sgRNAs and treated with DMSO or the USP1 inhibitor (USPi) ML-323 (30 μM) for 7 days. LE, long exposure; SE, short exposure. Phosphorylated (p)KAP1 on serine (S) 824 and γH2AX ubiquitylation (Ub) serve as markers of DSB signaling. **e,** Top: schematic of the S1 nuclease DNA fiber assay. Where indicated, cells were treated with ML-323 (30 μM). Bottom: measurements of IdU tract lengths in the indicated cells, with and without S1 nuclease treatment. A minimum of 100 tracts were measured per condition. Representative images are shown. **f,** Clonogenic survival assay of cells transduced with either a non-targeting or *RAD18*-targeting sgRNA. Cells were cultured for 10 days in the presence of DMSO, FEN1i (5 μM) and/or the USP1 inhibitor KSQ-4279 (2 μM). Data are representative of 2 experimental replicates. **g,** Competitive growth assay in cells expressing a *FEN1*-targeting sgRNA and the indicated cDNAs, in the presence of DMSO or KSQ-4279 (6 μM). cDNA expression was induced with 10 ng/ml doxycycline. 3 technical replicates are shown. **h,** A model depicting a potential mechanism underlying the *FEN1/LIG1*:*WDR48* synthetic lethal interactions. Gaps and unligated Okazaki fragments caused by LIG1 or FEN1 depletion trigger RAD18-dependent TLS. Dysregulation of this pathway by WDR48 loss/inhibition leads to persistent TLS, slow DNA synthesis, the accumulation of ssDNA gaps, DSBs, and cell death.

Second, to test the downstream involvement of TLS polymerases, we used the small molecule JH-RE-06 to prevent the interaction of REV1 with the REV7 subunit of Pol zeta, the major TLS extension polymerase^42,43^. JH-RE-06 efficiently rescued *LIG1*:*WDR48* synthetic lethality in a dose-dependent manner (Fig. 2b). To assess whether overactive TLS delays the completion of DNA replication in human cells, we synchronised cells in G0/G1 phase and monitored their progression through the cell cycle. In the presence of the USP1 inhibitor ML-323, LIG1-depleted cells progressed more slowly through S phase (Fig. 2c, Extended Data Fig. 3f, g). RAD18 depletion rescued this phenotype, with corresponding increases in G1 and G2/M phase cells.

Finally, we monitored DNA damage signaling and nascent DNA strand integrity. Prolonged treatment of LIG1-depleted cells with ML-323 led to higher levels of PCNA mono-ubiquitylation and a marked increase in DSB signaling, both of which were completely alleviated by RAD18 depletion (Fig. 2d). We speculated that these DSBs arise from post-replicative ssDNA gaps that are converted to DSBs in later S phases and are exacerbated by persistent TLS^44^. To test for post-replicative ssDNA gaps, we performed a DNA fiber assay coupled to S1 nuclease digestion (Fig. 2e)^45^. Control and LIG1-depleted cells were labelled with 5-chloro-2′-deoxyuridine (CldU) followed by 5-iodo-2′-deoxyuridine (IdU) in the presence or absence of ML-323. Permeabilised cells were incubated with S1 endonuclease which cleaves ssDNA gaps, generating DSBs and thereby shortening replication tract length. S1 nuclease treatment shortened the replication tracts of USP1-inhibited LIG1-depleted cells to a greater extent than all control cells (Fig. 2e). Thus, USP1 inhibition indeed leads to an accumulation of post-replicative ssDNA gaps in LIG1-depleted cells.

KSQ-4279 is a first-in-class USP1 inhibitor that potently and selectively kills HR-deficient cells and is currently in Phase 1 clinical trials in patients with HR-deficient advanced solid tumors^46^. We noted that *FEN1* is implicated as a tumor suppressor gene both in mouse models and human colorectal cancers^47,48^. We therefore asked if KSQ-4279 may have an unappreciated ability to preferentially kill FEN1-deficient cells. Clonogenic survival assays showed that FEN1i and KSQ-4279 co-treatment is synthetic lethal in a RAD18-dependent manner (Fig. 2f). Competitive growth assays demonstrated that FEN1-depleted cells were hypersensitive to KSQ-4279 treatment, and this was rescued by wild-type FEN1 cDNA expression (Fig. 2g, Extended Data Fig. 3h). These data suggest a potential use for USP1 inhibitors in the treatment of cancers with loss-of-function *FEN1* mutations^49^.

Overall, our data are consistent with a model in which WDR48-USP1 counteract hyperactive RAD18-mediated ubiquitylation of PCNA and subsequent TLS polymerase recruitment in FEN1/LIG1-depleted cells. Inappropriately high TLS at ssDNA gaps drives cell death in *FEN1*/*LIG1*:*WDR48* co-depleted cells (Fig. 2h). This function of WDR48-USP1 may also be important during unperturbed DNA replication, where a subset of Okazaki fragments evade repair by FEN1 and LIG1^21^. Our data further imply a use for USP1 inhibitors in tumors where ssDNA gaps accumulate such as *FEN1*-mutant colorectal cancers^48^.

### FANCM motor and SMARCAL1 annealing helicase activities ensure genome stability at late-replicating TA-rich repeats

We next turned to the interactions between *FANCM*/*FAAP24* and *SMARCAL1* as a second set of strong and novel genetic interactions identified in our screen (Fig. 1b, g, Extended Data Fig. 2j). FAAP24 and FANCM are well-established physical interaction partners with roles in ICL repair and the response to replication fork stalling^50,51^. FANCM is recruited to substrates by FAAP24, possesses ATP-dependent translocase activity and acts as a scaffold for numerous other DNA repair proteins^52^.

The SMARCAL1 ATPase facilitates the remodelling of stalled DNA replication forks, alongside factors such as ZRANB3 and HLTF^53,54^. Individual depletion of SMARCAL1, ZRANB3, or HLTF suppresses replication fork reversal, and these enzymes are thought to cooperatively regulate this process^55^.

Co-depletion of FANCM and SMARCAL1 profoundly reduced clonogenic survival (Fig. 3a, Extended Data Fig. 4a). We validated that combined loss of FANCM and SMARCAL1 is synthetic lethal using two different CRISPRi sgRNAs targeting each gene, as well as using CRISPRi knockdown of either *FANCM* or *SMARCAL1* in cells with isogenic knockout of the other partner (Extended Data Fig. 4b-d). The interaction was observed in p53-proficient RPE-1 cells as well as in HEK293 and K562 cells, demonstrating that it is independent of p53 status and not cell line specific (Extended Data Fig. 4e). Similarly, co-depletion of FAAP24 and SMARCAL1 was also synthetic lethal in individual assays (Extended Data Fig. 4f).

**Figure 3.**
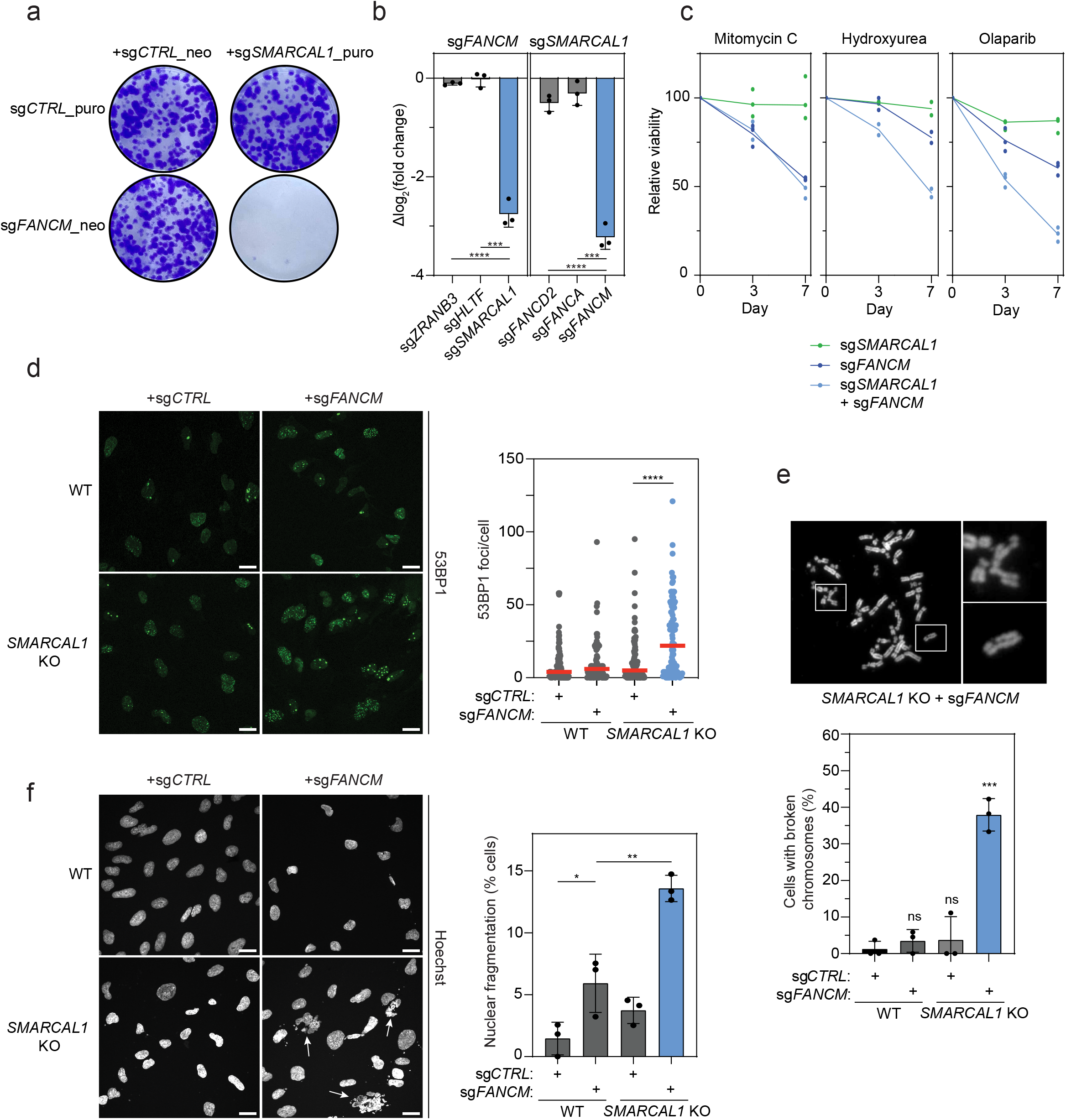
FANCM:SMARCAL1 loss leads to a defective replication stress response and genome instability: **a,** Co-depletion of FANCM and SMARCAL1 reduces clonogenic survival. RPE-1 *TP53* KO dCas9-KRAB were transduced with the lentiviruses containing sgRNAs, co-expressed with either a puromycin (puro)- or neomycin (neo)-resistance cassette, as indicated. Representative images of 10-day clonogenic survival are shown. Data are representative of 2 experimental replicates. **b,** Competitive growth assays for cells co-transduced with the indicated sgRNAs. The log_2_ fold change of co-depleted cells in a flow cytometry assay was normalized to the corresponding single depletions (sg*FANCM* or sg*SMARCAL1*). Values acquired on day 14 are shown. Error bars represent mean ± s.d; \*\*\**p* ≤ 0.001; \*\*\*\**p* ≤ 0.0001; unpaired Student’s *t* test. **c,** Dual-color flow cytometry survival curves for cells expressing the indicated sgRNAs and treated with mitomycin C (5 nM, 24h; *n = 3* technical replicates) hydroxyurea (2 mM, 24h; *n = 2* experimental replicates), or Olaparib (2 µM, chronic; *n = 3* technical replicates). Cell population ratios were normalized to those of corresponding untreated cells. A full stress sensitization panel can be found in Extended Data Fig. 5a. **d,** Left: representative images of 53BP1 foci in either WT or *SMARCAL1* KO cells transduced with a non-targeting or *FANCM-*targeting sgRNA. Right: quantification of 53BP1 focus formation. Red line denotes the median; ****p ≤ 0.0001; Mann-Whitney Test. A minimum of 80 cells derived from two experimental replicates were measured per condition. **e,** Top: a representative metaphase spread from *SMARCAL1* KO cells co-depleted of FANCM. Bottom: Quantification of metaphases with at least one broken chromosome in cells of the indicated genotype from three independent experiments. Error bars represent mean ± s.d; ns = not significant (*p* > 0.05); \*\*\**p* ≤ 0.001; unpaired Student’s t test. **f,** Left: representative images of nuclei in the indicated cell lines. Nuclei were stained with Hoechst. White arrows indicate cells exhibiting nuclear fragmentation. Right: quantification of nuclear fragmentation events from three experimental replicates. Error bars represent mean ± s.d; *p ≤ 0.05; **p ≤ 0.01; unpaired Student’s *t* test. Scale bars, 20 μm.

FANCM and SMARCAL1 are part of multi-component pathways, and we examined if each gene had synthetic lethal relationships with other components of these respective pathways. We observed no genetic interaction between *SMARCAL1* and FA genes involved in ICL repair and HR, such as *FANCD2* or *FANCA* (Fig. 3b). Surprisingly, *FANCM* deficiency was not synthetic lethal with loss of the other replication fork reversal enzymes *ZRANB3* or *HLTF*, despite efficient depletion of their transcripts (Fig. 3b & Extended Data Fig. 4g).

To delineate which functional domains of FANCM and SMARCAL1 are involved in their genetic interaction we performed dual-color flow cytometry assays in cells expressing cDNAs with defined point mutations or domain deletions (Extended Data Fig. 4h-k). Wild-type FANCM as well as variants lacking the FA core complex-binding MM1 (Δ943–1004) or BLM-TOP3A-RMI (BTR) complex-binding MM2 domain (Δ1219-1251) rescued the synthetic lethality. However, an ATPase-defective mutant (K117R) did not (Extended Data Fig. 4h, i). This indicates that the DNA-remodelling motor function of FANCM is required in SMARCAL1-deficient cells, but its recruitment of the FA and BTR complexes is dispensable.

Reciprocally, we tested whether SMARCAL1 variants lacking the N-terminal RPA-binding RBM (Δ2-32), the HARP2 domain required for DNA binding/annealing (W372A/F374A), or ATPase activity (R764Q and D549A/E550A) affected the synthetic lethal phenotype (Extended Data Fig. 4j, k). Expression of the ΔRBM, HARP2, and both ATPase-dead variants failed to rescue the viability, unlike wild-type SMARCAL1. This suggests that the abilities of SMARCAL1 to bind both RPA and DNA as well as hydrolyse ATP are required for cell survival in the absence of FANCM. The motor-driven annealing activity of SMARCAL1 at RPA-coated DNA could explain why other replication fork-remodelling translocases (ZRANB3 and HLTF) do not genetically interact with *FANCM*, since they do not contain RPA-binding domains.

Our data so far point to a new function of SMARCAL1 that is likely independent of the fork remodelling activity thought to be shared with *ZRANB3* and *HLTF*. To define this function, we sought to identify what type of lesion was driving the *FANCM*:*SMARCAL1* synthetic lethality. Using exogenous DNA damaging agents and measuring at early timepoints, we found that co-depleted cells were not further sensitised to ICLs (mitomycin C and formaldehyde) compared to FANCM depletion alone (Fig. 3c, Extended Data Fig. 5a). We observed no additive sensitivity to G4 quadruplex stabilisation (TMPyP4, PhenDC3, and Pyridostatin), replication fork stalling at common fragile sites (aphidicolin), or topoisomerase trapping (etoposide and camptothecin) (Extended Data Fig. 5a). Overexpression of RNase H1 failed to rescue the synthetic lethal phenotype, indicating that RNA-DNA hybrids are not involved (Extended Data Fig. 5b). However, *FANCM:SMARCAL1* co-depleted cells exhibited hypersensitivity to hydroxyurea and the PARP inhibitor, Olaparib, two agents that induce DNA replication stress and replication fork collapse through orthogonal mechanisms (Fig. 3a).

Replication fork collapse can lead to DSB formation. Indeed, even in the absence of exogenous DNA damaging agents, *SMARCAL1* knockout cells co-depleted for FANCM contained many more DSBs as quantified by 53BP1 foci (Fig. 3d). Metaphase spreads revealed that double-perturbed cells exhibited significantly elevated frequencies of broken chromosomes (Fig. 3e). These cells displayed high levels of nuclear fragmentation, a hallmark of mitotic DNA damage (Fig. 3f)^56^. Despite clear evidence of DSBs in *FANCM:SMARCAL1* co-deficient cells, they did not exhibit increased DSB signaling (Extended Data Fig. 5c). Consistent with previous reports, FANCM depletion alone resulted in diminished ATR signaling in response to hydroxyurea treatment, but this defect was not further affected by SMARCAL1 depletion^57^. These data suggest that simultaneous loss of FANCM and SMARCAL1 results in undetected DSB accumulation leading to chromosome instability after progression to mitosis.

To determine the physical context in which FANCM and SMARCAL1 are required to ensure genome stability, we performed MRE11 ChIP-Seq. This approach identifies recurrent sites of DSBs in wild-type, *SMARCAL1* knockout, FANCM-depleted, and double-perturbed cells (Fig. 4a)^58,59^. Relative to wild-type and single perturbation, double-perturbed cells exhibited very strong MRE11 binding at 72 unique genomic loci, 17 of which overlap known common fragile sites (Extended Data Fig. 6a)^60^. Motif analysis revealed a strong enrichment of interrupted TA-rich DNA sequences at these breakpoints (Fig. 4b-c, Extended Data Fig. 6b). These stereotyped TA repeats were very long, and much longer than unbroken TA repeats within the genome that are already prone to forming secondary structures such as hairpins (Fig. 4d)^61^. ChIP-qPCR revealed that FANCM physically binds one of these recurrently broken TA repeats in *SMARCAL1* knockout cells as compared with control cells or a non-TA-rich control locus (Extended Data Fig. 6c).

**Figure 4.**
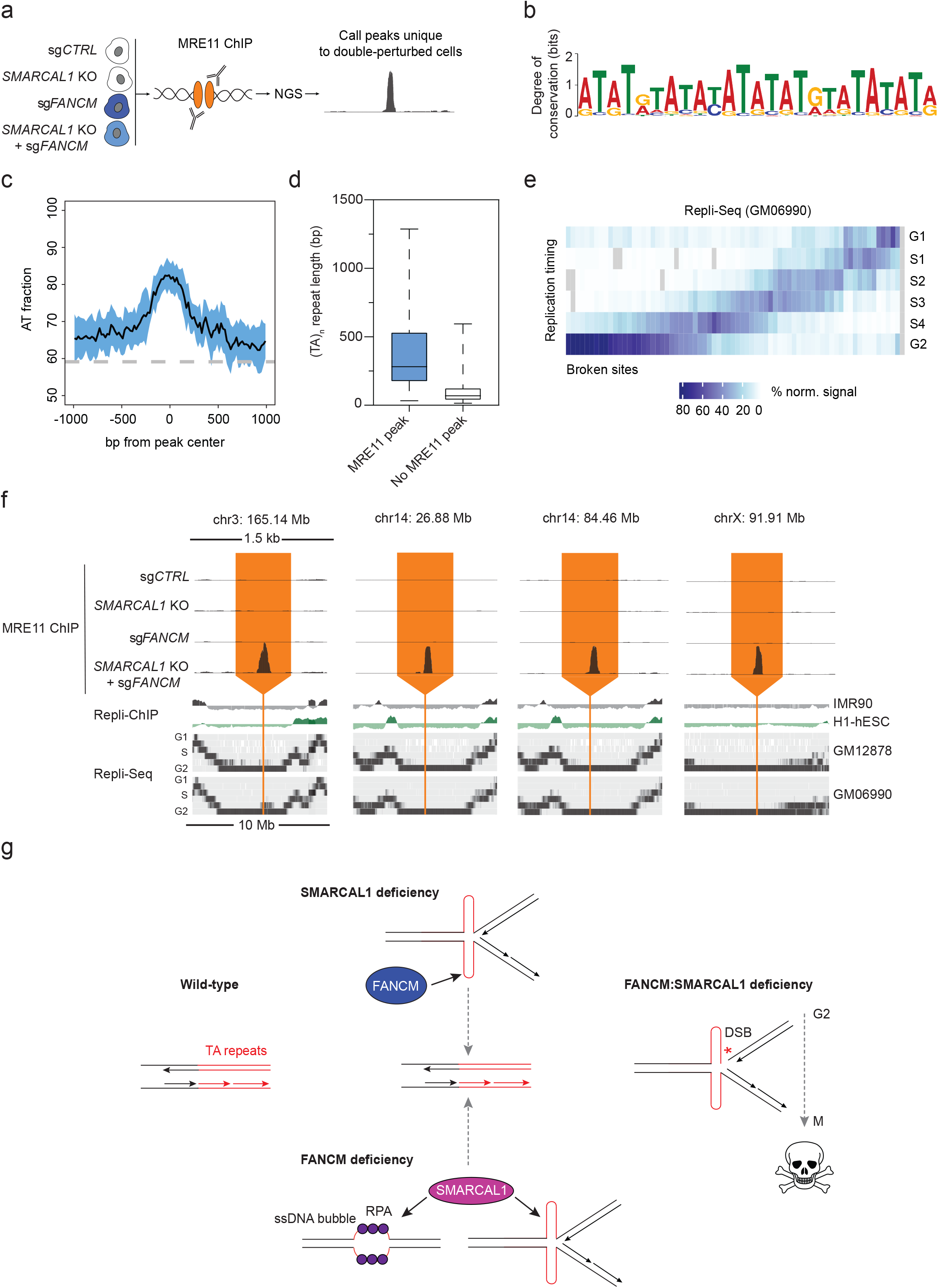
DSBs in FANCM:SMARCAL1-deficient cells occur at late-replicating TA-rich repeats: **a,** A schematic representation of the MRE11 ChIP-Seq approach used to identify sites of endogenous DSBs. 3 independent ChIP-Seq experiments were performed. **b,** Consensus motif identified by performing MEME analysis on DNA sequences underlying 72 MRE11 ChIP-Seq peaks only found in the *SMARCAL1* KO:sg*FANCM* cells. **c,** Plot showing the AT fraction per 20 base pair adjacent windows across 72 MRE11 ChIP-Seq peaks only found in the *SMARCAL1* KO:sg*FANCM* cells. Peaks were aligned according to their peak center. The AT fraction was calculated per 20 base pair in a 1 kb window. Blackline indicates the mean and blue area indicates standard deviation. The average AT fraction of the human genome is shown as a grey dashed line. **d,** Boxplot showing the lengths (1-99 percentile) of all annotated TA repeats in the hg19 reference genome, grouped according to whether they contain or are within 500 bp of an MRE11 ChIP-Seq peak exclusively detected in *SMARCAL1* KO:sg*FANCM* cells. The locations and lengths of (TA)_n_ repeats in the hg19 reference genome were taken from van Wietmarschen et al., 2020. **e,** Heatmap showing percentage-normalized signal from Repli-seq data in GM06990 cells across 72 MRE11 ChIP-Seq peaks only found in the *SMARCAL1* KO:sg*FANCM* cell. The average signal for each phase was taken per peak and ordered from late to early replication timing. **f,** Representative MRE11 ChIP-Seq tracks spanning a 1.5 kb window aligned with Repli-ChIP and Repli-seq tracks spanning a 10 Mb window in IMR90, H1-hESC, GM12878, and GM06990 cells. **g,** A model of the compensating activities of FANCM and SMARCAL1 in maintaining genome stability at TA repeats. SMARCAL1’s annealing motor activity could suppress secondary structure formation by favoring *trans* annealing of RPA-coated ssDNA. FANCM backs up this activity by unwinding secondary structures that do form. Lacking both SMARCAL1 and FANCM is particularly deleterious at late-replicating sites, where fork collapse leads to DSB formation upon entry into mitosis.

Strikingly, the breaks in *SMARCAL1* KO:sg*FANCM* cells were largely present in late replicating regions, with over half of the sites replicating in late S or even G2 (Fig. 4e, f, Extended Data Fig. 6d, e)^62^. Late formation of these DSBs potentially makes them less likely to be detected and repaired before cells enter mitosis. DSBs that persist into mitosis can lead to chromosome mis-segregation and cell death. Taken as a whole, our data suggest a model in which FANCM’s motor activity and SMARCAL1’s annealing motor activity prevent DNA breakage at late-replicating TA repeats during normal cell cycle progression (Fig. 4g).

## Discussion

Using a dual guide CRISPRi screen, we mapped the genetic interactions between 548 core DDR genes, encompassing all major DNA repair pathways and including essential genes. Our study led to new basic insight into the roles of important DNA repair factors. All the top synthetic lethal interactions were orthogonally validated, and the full genetic interaction network recapitulates known interactions and implicates new factors in diverse processes. We anticipate that our dataset will provide a platform for mechanistic investigations and drug target discovery. Mechanistically, we focused on two of the strongest synthetic lethal gene pairs identified in our screen: *FEN1*/*LIG1*:*WDR48* and *FANCM*:*SMARCAL1*.

We found that the USP1-activating function of WDR48 is essential to counteract RAD18 ubiquitin ligase activity and overactive TLS in cells where ssDNA gaps and unligated Okazaki fragments accumulate. We propose that DNA gaps trigger overactive TLS which drives cell death by slowing DNA synthesis, disrupting gap ligation, and exacerbating DSB formation. This aligns with data showing that TLS DNA polymerase kappa promotes DNA gap formation in BRCA1-deficient cells when USP1 is inhibited^44^.

Approximately 30-50 million Okazaki fragments are formed during each S phase and single-strand break repair proteins such as PARP1, LIG3, and XRCC1 are required to deal with those that inevitably evade canonical processing by FEN1 and LIG1^63,64^. Our work therefore implies a crucial role for WDR48-USP1 in suppressing overactive TLS at unligated Okazaki fragments during unperturbed DNA replication. The profound sensitivity of FEN1-deficient cells to the USP1 inhibitor, KSQ-4279, suggests it might be effective in treating *FEN1*-mutant cancers or augmenting chemotherapies that induce DNA gap formation, such as ATR or Wee1 inhibitors^48^.

Our identification of a genetic interaction between *FANCM* and *SMARCAL1* revealed previously undescribed roles for these factors in suppressing DSBs at long, late replicating TA-rich repeats. Long TA repeats can fold into secondary structures and stall DNA replication^61^. As an annealing helicase, SMARCAL1 efficiently brings together RPA-coated complementary ssDNA molecules in *trans*. We suggest that this activity suppresses secondary structure formation by preventing TA-rich repeats from self-annealing. Such a role could be important in other contexts. For example, SMARCAL1 occupies nucleosome-depleted regions near transcription start sites, which are also repetitive and prone to form secondary structures^65^. Alternatively, SMARCAL1 may be capable of directly unwinding secondary DNA structures, as it does R-loops^66^. In the absence of SMARCAL1, we propose that the translocase activity of FANCM unwinds secondary structures at TA repeats to facilitate DNA replication. When both SMARCAL1 and FANCM are absent, a failure to remove these roadblocks leads to DSB formation. But their very late replication timing limits DSB repair and mitotic catastrophe ensues.

*FANCM* is frequently mutated in multiple cancers, including triple-negative breast cancers, highlighting SMARCAL1 as a potential drug target^67^. FANCM may also represent a target of therapeutic intervention in cancers with loss-of-function *SMARCAL1* mutations, which are prevalent in glioblastomas^68^. Our DDR interaction map identifies numerous other synthetic lethal relationships that could be exploited in cancer therapy. Many of these relationships already link established cancer genes and existing small molecule targets and thus are immediately actionable. We anticipate that exploration of our dataset will define additional cancer-specific vulnerabilities that could translate into clinical benefit in patients.

## Materials & methods

### Cell culture

hTERT RPE-1 and HEK293(T) cell lines were cultured in Dulbecco’s Modified Eagle’s Medium (DMEM/F12; Merck) supplemented with 10% fetal bovine serum (FBS; Thermo Fisher Scientific) and 100 units/mL streptomycin and 100 mg/mL penicillin (GIBCO). K562 cells were cultured in RPMI-1640 with 25mM HEPES and 2.0 g/L NaHCo3 in 10% FBS, 2 mM glutamine, and 100 units/mL streptomycin and 100 mg/mL penicillin. All cell lines were grown under 3% oxygen. Cells were routinely tested for mycoplasma.

### Cell line generation

hTERT RPE-1 *TP53* KO cells were obtained from the Stephen Jackson Lab (Cambridge). RPE-1 *TP53* KO dCas9-KRAB cells were generated by transduction with a lentiviral vector encoding the dCas9-KRAB and a Blasticidin resistance cassette with a low MOI. Cells were selected with Blasticidin, and single cell clones were seeded by cell sorting. Resulting clones were validated for CRISPRi activity by CD55 knockdown efficiency. RPE-1 *TP53* KO *FANCM* KO and *SMARCAL1* KO cell lines were generated using CRISPR/Cas9. sgRNAs were selected from the VBC website and were in vitro transcribed with T7 RNA polymerase^69^. Editing of cells was done as described previously^70^. A LONZA AMAXA nucleofector was used with P3 solution and program EA104. Clones were single cell seeded by serial dilution and clones were selected using Sanger Sequencing and ICE analysis (Synthego). Knockout of FANCM and SMARCAL1 was validated by NGS and Western blotting.

### Cloning and mutagenesis

For RNaseH1 WT or WKKD mutant (W43A, K59A, K60A), cDNAs from Addgene plasmids #111906 and #111905 were PCR amplified. Resulting fragments and mCherry cDNA amplified from Addgene plasmid #102245 were assembled using Gibson Assembly. Empty and FANCM (WT and K117R) pLVX-TetOne-Puro plasmids were a generous gift from the Claus M. Azzalin lab. Point mutants were created by site-directed mutagenesis. Internal deletion mutants were generated by inverse PCR, followed by T4 PNK phosphorylation and blunt-end ligation. 3xFLAG-SMARCAL1 cDNAs were cloned into the lentiviral vector pLVX-TetOne-Puro (Clontech) by Gibson Assembly. For all Gibson Assemblies the NEB Hi-Fi Assembly Mastermix (NEB) was used. All PCR reactions were performed using Q5 DNA polymerase.

### sgRNA cloning

CRISPRi sgRNAs were cloned into either BFP- or GFP-containing lentiviral vectors (Addgene plasmids #60955 and #111596). To generate double knockdown cells sgRNAs were ligated into IGI-P1589 containing a Neomycin resistance cassette. Forward and reverse primers for each sgRNA were annealed by pre-incubation at 37°C for 30 minutes with T4 polynucleotide kinase (PNK; NEB), followed by incubation at 95°C for 5 minutes and then ramp down to 25°C at 5°C/min. Annealed sgRNAs were ligated into the corresponding vector that had been digested with BstXI and BlpI restriction enzymes using T4 Ligase (NEB). All sgRNA protospacers are listed in Supplementary Table 3.

### Design of double-sgRNA CRISPRi SPIDR library

Genes to target in the SPIDR sgRNA library (n=548) were selected by all genes with the gene ontology term “DNA repair” (GO:0006281). For each gene, sgRNAs from the human CRISPRi-v2 library were ranked using a strategy described previously and using a preliminary dataset of >50 CRISPRi screens^14,16^. Briefly, genes were divided into three tiers. For genes essential for growth in K562 cells (tier 1), sgRNAs were ranked by their growth phenotypes. For genes that had scored as a significant hit in at least 4 CRISPRi screens (tier 2), sgRNAs were ranked by their average phenotype across all screens in which the gene had scored as a hit. For all other genes (tier 3), sgRNAs were ranked by the regression scores from the human CRISPRi v2.1 algorithm.1 For each gene, two sgRNAs were then selected for each transcription start site targeted in the human CRISPRi-v2 library, as follows:

For genes (n=249) for which knockout caused strong growth phenotypes in either RPE-1 cells or in at least 5 neuroblastoma cell lines, as assessed by querying data from 17 screens in neuroblastoma cell lines (identifiers NB5, NB6, NB7, NB10, NB13, NB17, NB69, CHP-212, SK-N-FI, SK-N-AS, SK-N-DZ, SK-N-SH, KP-N-YN, KP-N-YS, TGW, BE2-M17, NH-12) in the Project Score database from the Cancer Dependency Map (https://score.depmap.sanger.ac.uk/, accessed August 2020), these two sgRNAs were: 1) the top sgRNA identified using the algorithm above, and 2) a mismatched variant of the top sgRNA, chosen to have empirical relative activity of 0.47 ± 0.15 if an sgRNA existed for which relative activity in this window had been measured or otherwise chosen as the mismatched variant with predicted activity closest to 0.5 out of all possible singly mismatched variants, with activity predictions derived from a convolutional neural network as previously described^15,71,72^. Mismatched sgRNAs were required to have an on-target specificity score of at least 0.15, calculated as described previously^15^.

For genes (n=299) for which knockout did not cause strong growth phenotypes in both RPE-1 and >12 neuroblastoma cell lines, the top two sgRNAs from the algorithm above were selected.

From this set of sgRNAs, the double-sgRNA libraries were assembled in a programmed manner. Each unique combination of two genes was randomly assigned an orientation of “ab” or “ba”. For each such combination, all sgRNAs targeting the gene in position a were paired with all sgRNAs targeting the gene in position b. Each sgRNA targeting a given gene was also paired with all other sgRNAs targeting that gene. Each unique sgRNA in the library was furthermore paired with 15 different non-targeting negative control sgRNAs in both the ab and the ba orientations. Finally, 225 pairs consisting of all possible combinations of 15 different non-targeting negative control sgRNAs were included as negative controls.

All sgRNA pairs were then ordered as an oligo pool from Agilent Technologies, containing the following constant sequences: AACTGCGATCGCTAATGTCCACCTTGTTG (upstream of sgRNA a), gtttcagagcgagacgtgcctgcaggatacgtctcagaaacatg (between sgRNA a and sgRNA b), GTTTAAGAGCTAAGCTGGTTCTCCAGTGCCTTATT (downstream of sgRNA b).

### Dual guide library cloning

Dual guide library cloning was done as described in Replogle et al., 2022 with small modifications. After PCR amplification, oligos were BstXI/BlpI-digested and ran on a polyacrylamide gel. The corresponding band was extracted, and DNA was purified before ligation into pJR103 (Addgene plasmid #187242). The ligated DNA was precipitated using isopropanol (Thermo Fisher Scientific, Cat# 17170576) and transformed into Mega-X electrocompetent cells (Thermo Fisher Scientific, Cat# C640003). Bacteria were cultured in lysogeny broth overnight and plasmid was harvested with Plasmid Plus Midi Kit (QIAGEN). This intermediate library and pJR98 were digested using BsmBI and the insert from pJR98 was ligated into the intermediate library to insert constant region CR3* for the first sgRNA and the hU6 promoter for the second sgRNA. A library coverage of at least 30x was continually maintained during the cloning process. The final library expresses the first sgRNA under the mU6 promoter with CR3*, while the second sgRNA is expressed under a hU6 promoter with CR1.

### Lentivirus packaging

Lentiviruses were produced in HEK293T cells. Briefly, HEK293T cells were transfected using polyethylenimine (PEI) with the VSV-G envelope expressing plasmid, pMD2.G, the packaging plasmid, psPAX2, and our transfer vector. Lentiviral supernatant was harvested 48- and 72-hours post-transfection.

### CRISPRi screen

hTERT RPE-1 *TP53* KO cells stably expressing dCas9-KRAB were transduced with the lentiviral library at a MOI of ∼0.25 with a coverage of at least 350 cells per sgRNA in media supplemented with polybrene (10 μg/ml). The next day, the culture media was replaced with puromycin-containing media (15 μg/ml) to select for transductants. Selection was carried out for 96 hours, during which time cells were expanded and divided into 2 technical replicates. Hereafter, library coverage was maintained at ∼500 cells per sgRNA combination. 4 days after transduction, the background/timepoint 0 samples were harvested and cell pellets were frozen (-80°C). The remaining cells were seeded and continually sub-cultured when near 100% confluency for 14 days (∼10 population doublings), at which time cell pellets were harvested and frozen (-80°C). Genomic DNA (gDNA) was isolated using the Gentra Puregene Cell Kit (QIAGEN). Integrated sgRNA regions were amplified by PCR using custom primers (Supplementary Table 4) and paired-end sequencing was performed on a NovaSeq 6000.

### Screen analysis

Raw FASTQ files were processed using seal (BBMap – Bushnell B. – sourceforge.net/projects/bbmap/) to search for sgRNA matches in the first 22bp of each read with hamming distance of 0 and a kmer length of 20bp. A custom python script was used to parse the annotated FASTQ files and generate a count matrix containing all possible sgRNA pairs. The coverage per sgRNA pair was calculated to assess the quality of each sample (Extended Data Fig. 1a-b). Total counts were normalized using the median ratio method to estimate size factors using the non-targeting sgRNAs^73^. sgRNA pairs with <10 read counts after normalization were filtered out. R version 4.1.2 was used to generate custom plots.

Normalized counts were used to obtain LFCs from GEMINI analysis. Count data were filtered with having a representation ≥50 reads in the T0 sample of each replicate. LFCs were calculated using the GEMINI package with a pseudo-count of 10 and a modified gemini_calculate_lfc_mod function to prevent double-normalization (see code availability). Essential sgRNAs were determined with having an average LFC of < -3 at T14 when combined with non-targeting sgRNAs. Perfect and mismatched variants of these sgRNAs were included as separate genes in the analysis for Extended Data Fig. 1f to determine recovery of genetic interactions for strong essential genes. Mismatched variants of strong essential sgRNAs showed a reduced signal at T14 and were able to recover additional genetic interactions (Extended Data Fig. 1e-f). Therefore, the count data were filtered for these candidates and only the mismatched sgRNA variant was kept for all analysis. Non-targeting sgRNAs were used as the negative control gene to build the GEMINI model. Sensitive and strong GEMINI scores were calculated with a λ of 1 to identify synthetic lethal interactions of a broad range of genes. To identify sensitive viable (recovery) interactions in Fig. 1b, a λ of 0.5 was used. GEMINI scores > 0 were considered potential genetic interactions and all combinations with a score < 0, were considered non-interacting and a score of 0 was assigned. UMAP embedding was calculated in R using the strong score across a symmetric version of the entire dataset without prior dimension or variance reduction.

### Dual-color flow cytometry assay

The indicated cells were transduced with sgRNA-containing lentiviruses at the same MOI. After 72 hours, cells were seeded in triplicate in 6-well plates and, after an additional 24 hours, cell populations were analysed by flow cytometry (= day 0) using an Attune NxT Flow Cytometer (Invitrogen). Cell populations were analysed at regular time intervals (as indicated in the Figures) thereafter. For triple knockdown and cDNA experiments, cells were transduced with the relevant sgRNA-containing lentiviruses and selected with puromycin for 96 hours prior to being transduced with lentiviruses containing sgRNAs targeting synthetic lethal gene pairs. For cDNA experiments, expression was induced with doxycycline 24 hours before transduction with lentiviruses containing *SMARCAL1* and *FANCM-*targeting sgRNAs. Doxycycline-containing medium was refreshed every 48 hours. Where drug treatments were performed, cells were treated 24 hours after seeding on 6-well plates (on day 0).

### Clonogenic survival assays

hTERT RPE-1 *TP53* KO cells were co-transduced with lentiviruses containing sgRNAs that were co-expressed with either a puromycin- or neomycin-resistance cassette. After selection with puromycin (10 μg/ml) and/or G418 (1.25 mg/ml), 1000 cells of each sample were seeded in triplicate on 6-well plates. Where indicated, drug treatments were performed 24 hours after seeding. 10 days later, cells were rinsed in PBS and stained with 0.5% (w/v) crystal violet in 20% (v/v) methanol for 15 minutes at room temperature. After staining, plates were rinsed with H_2_O and air-dried.

### Competitive growth assays

Cells were independently transduced with either a non-targeting sgRNA (co-expressed with either mCherry, GFP, or BFP) or a targeting sgRNA. After 24 hours, cells were treated with the appropriate antibiotic - either puromycin (10 μg/ml) or G418 (1.25 mg/ml) - until untransduced cells were selected out of the population. Cells were then mixed at a 1:1 ratio by number and seeded in triplicate on 6-well plates. After 24 hours, cell populations were analysed by flow cytometry. Cells were treated or not with exogenous agents as indicated in the figures and monitored by flow cytometry at the indicated timepoints thereafter.

### Drug treatments

The chemicals used in this study can be found in Supplementary Table 4. Treatment durations and doses are indicated in the figures/legends.

### Quantitative reverse transcription (RT)-qPCR

1,000,000 RPE-1 cells were harvested, and total RNA was isolated using the RNeasy Kits (QIAGEN), according to the manufacturer’s instructions. RNA was reverse transcribed into cDNA using iScript™ Reverse Transcription Supermix (Bio-Rad) using oligo dT primers. qPCR reactions were prepared with the SsoAdvanced Universal SYBR Green Supermix (Bio-Rad) and ran on a QuantStudio 6 Flex Real-Time PCR System (Applied Biosystems). Primers used for qPCR can be found in Supplementary Table 4. Data were analyzed using the Delta Delta Ct (ΔΔCt) method.

### Nucleofection of RNaseH1 constructs

For transient expression of RNaseH1 constructs, 100,000 RPE-1 cells were transfected with 1 μg of RNaseH1 plasmid using the LONZA AMAXA nucleofector with P3 solution and program EA104. Nucleofected cells were recovered in warm media and identified by mCherry expression by fluorescence-activated cell sorting (FACS).

### Cell cycle analysis

hTERT RPE-1 *TP53* KO cells were arrested in G0/G1 by contact inhibition for 48 hours then re-plated at low density (100,000 cells/well on a 6 well plate) to release them into S phase in the presence or absence of ML-323 (30 μM). Cells were harvested at the indicated timepoints, fixed in 70% ethanol for 30 minutes on ice, washed once with staining buffer (1X PBS + 5% FBS), and stained at room temperature using 1 µg/ml DAPI solution (BD Biosciences) diluted in staining buffer for 5 minutes and analysed using an Attune NxT Flow Cytometer. Subsequent analysis was performed using FlowJo v10.8.1.

### Western blotting

Cells were lysed in radioimmunoprecipitation assay (RIPA) buffer (0.5M Tris-HCl, pH 7.4, 1.5M NaCl, 2.5% deoxycholic acid, 10% NP-40, 10mM EDTA), supplemented with Halt Protease Inhibitor Cocktail and Phosphatase Inhibitor Cocktail (ThermoFisher). Samples were sonicated using a Bioruptor Plus sonicator (30 secs ON, 30 sec OFF for 5 cycles) and centrifuged for 5 minutes at 21,000 x g. Protein concentrations were measured using a Bradford assay. Samples were normalized by protein concentration, mixed with 4x NuPage LDS sample buffer supplemented with 5% β-mercaptoethanol, and boiled for 5 minutes at 98°C. Samples were loaded into NuPAGE™ 4-12% Bis-Tris Protein Gels (ThermoFisher) and transferred to 0.2 μm nitrocellulose membranes. Membrane blocking was performed with 5% milk diluted in Tris-buffered saline containing 0.1% Tween 20 (TBS-T). Primary antibodies were diluted in 5% BSA diluted in TBS-T. After incubation with primary antibodies, membranes were incubated with Li-Cor near-infrared fluorescence secondary antibodies, after which they were scanned using a Li-Cor Near-InfraRed fluorescence Odyssey CLx Imaging System. All antibodies used are listed in Supplementary Table 4.

### Immunohistochemistry

Cells were seeded 48 hours before on sterile 15 mm glass coverslips. Coverslips were washed in ice cold PBS and nuclei were 5 minutes pre-extracted by incubating cells in pre-extraction buffer (HEPES pH 7.5 (25 mM), NaCl (50 mM), EDTA (1 mM), MgCl_2_ (3 mM), Sucrose (300 mM), Triton X-100 (0.5%)) on ice. Cells were washed once in PBS and then fixed in 4% formaldehyde for 15 minutes at RT. Coverslip were again washed two times in PBS before blocking in 5% BSA in PBS ON at 4°C. Primary antibody incubation with 53BP1 antibody (NB100-304) was carried out ON at 4°C. After three washes in PBS, coverslips were incubated with the secondary Alexa Fluor™ 488 antibody for 1 hour at RT. Coverslips were washed three times in PBS and then incubated with Hoechst 33342 Ready Flow™ Reagent for 5 minutes before two additional wash steps in PBS and a Milli-Q water wash. Cells were mounted onto glass slides using ProLong™ Diamond Antifade Mountant. All antibodies were diluted in 2.5% BSA in PBS. All incubation steps were carried out in a wet chamber. Images were taken using Leica SP8 confocal microscope or ZEISS Apotome 3. For foci analysis, the layer with the highest intensity was selected. If there were foci in multiple planes a z-stack and maximum intensity projection was performed. For quantification of mitotic catastrophes, Hoechst staining was used to detect nuclear morphology and fragmented nuclei were scored as shown in Fig. 3f. The data were quantified using Fiji-ImageJ 2.9.0.

### Metaphase spreads

Cells were treated with 0.04 μg/ml colcemid for 16 hrs to arrest them in mitosis then collected, washed with PBS, and incubated in 0.075 M KCl at 37°C for 10 minutes. Cell pellets were fixed in methanol:acetic acid (3:1 ratio), spread on glass slides, then coated with ProLong™ Diamond Antifade Mountant. Spreads were imaged using a ZEISS Apotome 3 microscope.

### S1 nuclease DNA fiber assays

The S1 nuclease DNA fiber assay was conducted as described in Jacobs et al., 2022, with the following modifications. Cells were incubated with 19 mM CldU for 15 minutes, washed three times with warm PBS, then incubated with 28 mM IdU for 45 minutes. During the IdU incubation, cells were treated or not with ML-323 (30 μM). Cells were then harvested and incubated with CSK-100 buffer (100 mM NaCl; 20 mM HEPES; 3 mM MgCl_2_; 300 mM sucrose; 0.5% Triton X-100) for 5 minutes, then washed with PBS. Cells were then incubated in 200μL S1 nuclease reaction buffer with or without S1 nuclease (20 U/mL; ThermoFischer Scientific) for 30 minutes at 37°C. Cells were then washed with PBS, lysed on glass slides in a buffer composed of 200 mM Tris-HCl, pH 7.5, 50 mM EDTA, and 0.5% SDS for 5 minutes, and spread by tilting the slides at a 45° angle. The slides were then air dried and fixed in methanol/acetic acid (3:1 ratio) overnight at 4°C. The slides were then incubated in 2.5 M HCl for 1 hour to denature the DNA fibers, washed with PBS and incubated in 1% BSA/PBS (0.2% Tween 20) for 40 minutes. Staining was performed by incubating the slides for 2.5 hours at room temperature with anti-CldU (1:500, ab6326; Abcam) and IdU (1:100, B44, 347,580; BD Biosciences) antibodies, followed by 1 hour room temperature incubation with anti–mouse Alexa Fluor 488 (1:300) and anti–rat Alexa Fluor 568 (1:150) secondary antibodies. All antibodies were diluted in 2.5% BSA/PBS. Fibers were visualized using a Leica SP8 confocal microscope or ZEISS Apotome 3 microscope (64x, oil) and analysed using Fiji-ImageJ 2.9.0.

### ChIP-Seq

Ten million cells for each condition were fixed in 1% formaldehyde at room temperature for 15 minutes. The fixation reaction was quenched with glycine to a final concentration of 125 mM. Cells were harvested and washed twice with chilled PBS, and pellets were snap frozen in a dry ice-ethanol bath before storing at -80°C. When ready to process, cell pellets were thawed on ice and incubated with lysis buffer (LB) 1 (50 mM HEPES–KOH, pH 7.5; 140 mM NaCl; 1 mM EDTA; 10% Glycerol; 0.5% NP-40 or Igepal CA-630; 0.25% Triton X-100; 1X protease inhibitors) on ice for 10 minutes. Cells were pelleted by centrifugation and incubated with LB2 (10 mM Tris–HCL, pH8.0; 200 mM NaCl; 1 mM EDTA; 0.5 mM EDTA; 1X protease inhibitors) for 5 minutes on ice. The extracted nuclei were pelleted by centrifugation and resuspended in LB3 (10 mM Tris–HCl, pH 8; 100 mM NaCl; 1 mM EDTA; 0.1% NaDeoxycholate; 0.5% N-lauroylsarcosine; 1X protease inhibitors). Nuclei were sonicated using a Covaris S2 sonicator with the following settings: duty cycle 5%, intensity 5, 200 cycles per burst, 12 minutes. Debris was pelleted by centrifugation at 4°C and the supernatant was transferred to a 5 mL tube. 100 µL of Dynabeads protein A that had been prebound with MRE11 antibody (Novus, Cat# NB100-142) were added to the cell lysate and samples were incubated at 4°C overnight with rotation. Beads were collected on a magnetic stand and washed with ice-cold RIPA buffer 6 times, followed by a final wash with TBS before resuspending beads in 200 µL of elution buffer (50 mM Tris–HCl, pH 8; 10 mM EDTA; 1% SDS). The bead slurries were incubated overnight at 65°C to reverse crosslinks. Samples were treated with 1 mg/mL RNaseA (Ambion, Cat# 2271) for 30 minutes at 37°C, followed by proteinase K treatment 20 mg/mL (Invitrogen, Cat# 25530-049) for 1 hour at 55°C. DNA was then purified using a MinElute PCR Purification Kit (Qiagen, Cat# 28004) and sequencing libraries were prepared using a NEBNext Ultra II kit.

### ChIP-Seq analysis

Raw paired-end FASTQ files were aligned to the GRCh38 reference genome using bowtie2 version 2.4.4. Resulting BAM files were sorted and indexed with samtools version 1.6. Peaks were called using Macs3 version 3.0.0b1 with RPE1 WT set as the control and default settings. Peaks were filtered using the ENCODE blacklist v2 and manually inspected^74^. Total read counts were calculated for each sample and used to generate comparable bigwig files for visualization using bamCoverage version 3.3.0 and the scaleFactor option. CrossMap version 0.6.0 was used to lift over GRCh38 to GRCh19 coordinates. Locally hosted bigwig files were added as custom tracks to the UCSC Genome Browser for visual inspection. MEME-ChIP version 5.5.1 was used to identify motifs around the peaks and ChIPseeker was used to visualize peak locations over chromosomes. The AT fraction was calculated per base across all peaks in a 1000bp window around the center of each peak.

### Repli-Seq Analysis

Repli-Seq data for GM06990 cells from the UW ENCODE group were downloaded from the UCSC genome browser as bigwig files in wavelet-smoothed and percentage-normalized signal format (GEO accession: GSM923443). Deeptools version 3.5.1 was used to visualize the wavelet-smoothed signal data for all peaks in a 2 Mb region around each peak center with the tools computeMatrix reference-point, with the bin-size option set to 1, and plotHeatmap to create the heatmap. R was used to process and plot the percentage-normalized signal data.

### V5-FANCM ChIP-qPCR

Expression of V5-tagged FANCM was induced with 1 µg/ml doxycycline in wild-type or *SMARCAL1* KO hTERT RPE-1 *TP53* KO cells 48 hours prior to harvesting. ChIP-qPCR was performed largely as described previously, with the following modifications^52^. Chromatin shearing to ∼500-1500 bp was performed using a Diagenode Bioruptor 300 for 40 cycles (30 seconds ON; 30 seconds OFF). Protein G coupled magnetic dynabeads were pre-coated with 5 µg anti-V5 antibody for 4 hours at 4°C. DNA concentrations were measured on a TapeStation high-sensitivity flow cell (Agilent). qPCR reactions were performed with SsoAdvanced Universal SYBR Green Supermix (Bio-Rad) and analysed on a QuantStudio 6 Flex Real-Time PCR System (Applied Biosystems). Primers used for qPCR can be found in Supplementary Table 4.

## Supplementary figure legends

**Extended Data Figure 1 – related to Figure 1. Quality control of the screen and analysis of mismatched guide RNAs:** Quantification of log_2_ pair count of replicate 1 at **a,** T0 and **b,** T14 shows high coverage at T0 and drop out of many combinations at T14. **c,** Comparison between screen replicates at T0 and T14. **d,** Volcano plot of log_2_ fold change of non-targeting (sg1 & sg2 non-targeting) and single gene (sg1 == sg2) targeting library elements at T14 compared to T0. Common essential genes (e.g., *MCM2*, *GTF2H2*) show high negative values while tumor suppressors *SAMHD1* and *TAOK1* show a positive log_2_ fold change, respectively. **e,** Growth phenotypes of mismatched sgRNAs for strongly essential genes. Violin plot showing the log_2_ fold change of essential sgRNAs (LFC ≤ -3) and their mismatched variant. Mismatched variants have significantly decreased growth phenotype, \*\*\*\**p* ≤ 0.0001; Wilcoxon rank sum test. Density plot for essential sgRNAs showing delta LFC for mismatched minus perfect sgRNAs. Most mismatched variants reduce the negative growth phenotype, leading to the majority of delta LFC density > 0. Vertical dashed line indicates delta LFC of 0. **f,** Sensitive GEMINI score rank-ordering of all binary interactions with perfect sgRNAs or mismatched sgRNAs (asterisk) for essential genes. Mismatch sgRNAs help to recover known (magenta) as well as novel (light blue) genetic interactions. Mismatched guides were used for strongly essential genes for all further analysis. **g,** Boxplots showing the sensitive GEMINI scores for cells expressing the indicated sgRNAs.

**Extended Data Figure 2 – related to Figure 1. Orthogonal validation of novel synthetic lethal gene pairs: a,** Left: Schematic representation of the dual-color flow cytometry assay. Cells are transduced with sgRNAs and assayed 96 hours later and at regular intervals thereafter. Right: An example of the flow cytometry gating strategy. **b,** Representative flow cytometry density plots of the *FEN1:WDR48* interaction in RPE-1 *TP53* KO dCas9-KRAB cells. **c-i,** RPE-1 *TP53* KO dCas9-KRAB cells were co-transduced with the indicated sgRNAs. Cells populations were first measured by flow cytometry 96 hours after transduction (day 0) and on the indicated days thereafter. Log_2_ fold changes relative to day 0 are shown and all values were normalized to the corresponding untransduced control cells. These panels show the full quantifications of data plotted in Fig. 1e. **j,** Stringently filtering for only the strongest synthetic relationships (GEMINI sensitive score ≤ -2.5) reveals two unanticipated synthetic lethal modules: *WDR48* (also known as *UAF1*) with *LIG1* and *FEN1*, and *SMARCAL1* with *FANCM* and *C19orf40/FAAP24*.

**Extended Data 3 – related to Figure 2. WDR48 is required for the survival of cells lacking single-strand break repair/Okazaki fragment maturation factors: a,** All genetic interactions of *LIG1* identified in the SPIDR screen. **b,** Flow cytometric quantifications of the genetic interactions between *FEN1* and the genes encoding the WDR48-interacting proteins USP1, USP12, and USP46. The log_2_ fold change of the co-depleted cells was normalized to the corresponding single depletions. Values acquired on day 14 are shown. **c,** Western blot of control or WDR48-depleted cells RPE-1 *TP53* KO dCas9-KRAB cells treated with DMSO or FEN1-IN-1 (FEN1i; 10 μM) for 7 days. **d,** Quantification of competitive growth assays with cells transduced with the indicated sgRNAs. Cell populations were monitored by flow cytometry after a 10-day treatment with DMSO or FEN1i (10 μM). **e,** Competitive growth assays for cells expressing the indicated sgRNAs and treated with DMSO or ML-323 (30 μM). Cell population ratios were monitored at the indicated time intervals by flow cytometry and normalized to the corresponding untreated controls. **f,** Quantification of Fig. 2c. Error bars represent mean ± s.d. \*\*\**p* ≤ 0.001; \**p* ≤ 0.05; ns = not significant (*p* > 0.05); paired Student’s *t* test. **g,** Quantification of all time points for experiments in Fig. 2c. **h,** Western blot validating FEN1 knockdown and cDNA expression for experiments shown in Fig. 2g. 3 technical replicates are shown for b, d, e, f, and g.

**Extended Data 4 – related to Figure 3**. **Validation and characterization of the *FANCM*:*SMARCAL1* synthetic lethal interaction: a,** Western blot validating sgRNA-mediated knockdown of *FANCM* and *SMARCAL1*. **b,** Quantifications of the dual-color flow cytometry assay measurements of cells transduced with the indicated sgRNAs. Values were normalized to the log_2_ fold change of the corresponding untransduced cells. The log_2_ fold change on day 14 is shown. **c,** Western blot of whole cell extracts of *FANCM* and *SMARCAL1* knockout (KO) cells. The asterisk denotes a non-specific band. **d,** Competitive growth assays of wild-type (WT) and *FANCM* or *SMARCAL1* CRISPR knockout RPE1 *TP53* KO dCas9-KRAB cells transduced with the indicated sgRNAs with the same MOI. Values were normalized to day 0 and the relative log_2_ fold changes are shown. **e,** Dual-color survival curves of dCas9-KRAB-expressing RPE1 *TP53*-proficient, HEK293, and K562 cells transduced with the indicated sgRNAs. Values were normalized to the corresponding untransduced population of cells and log_2_ fold changes relative to day 0 are shown. **f,** Quantification of the dual-color FACS assay for cells expressing sgRNAs targeting *FAAP24* and/or *SMARCAL1.* Experiments were performed as in Extended Data Fig. 4e. **g,** Validation of CRISPRi-mediated knockdown of *ZRANB3* and *HLTF* by quantitative RT-PCR (RT-qPCR). mRNA expression was normalized to the sgCTRL sample. 3 technical replicates are shown. **h,** Quantification of the dual-color flow cytometry assay for FANCM:SMARCAL1 co-depleted cells complemented with the indicated FANCM cDNAs. cDNA expression was induced with 1 µg/ml doxycycline. The log_2_ fold change was normalized to Day 0. **i,** Western blot controls of cDNA expression for Extended Data Fig. 4h. **j,** Dual-color flow cytometry assay for FANCM:SMARCAL1 co-depleted cells complemented with the indicated SMARCAL1 cDNAs. Experiments were performed as in Extended Data Fig. 4h, except cDNA expression was induced with 0.25 µg/ml doxycycline. **k,** Western blot controls of cDNA expression for Extended Data Fig. 4j. \**p* ≤ 0.05; \*\**p* ≤ 0.01; \*\*\*\**p* ≤ 0.0001; ns = not significant (*p* > 0.05); Unpaired Student’s *t* test that compared cells expressing WT cDNAs with each other condition on day 14. 3 technical replicates are shown for b, e, f, h, and j.

**Extended Data 5 – related to Figure 3. Further characterization of the *FANCM*:*SMARCAL1* synthetic lethal interaction: a,** Drug response assays of FANCM, SMARCAL1, and co-depleted cells. Cells were treated with genotoxic agents (APH, aphidicolin; PDS, pyridostatin; ETO, etoposide; CPT, camptothecin) 96 hrs after transduction with the indicated sgRNAs. Concentrations and treatment durations are indicated above each panel. Cells were re-analysed by FACS 7 days later. Cell population ratios were normalized to those of corresponding untreated cells. 3 technical replicates are shown. **b,** Survival curves of RPE1 *TP53* KO cells transduced with sgRNAs targeting FANCM and SMARCAL1 and nucleofected with a cDNA encoding either wild-type (WT) or R-loop-binding defective (WKKD) RNase H1. Cell populations were measured by flow cytometry and normalized to corresponding untransduced control cells. **c,** Western blot of whole cells extracts from cells transduced with the indicated sgRNAs and either left untreated or treated with 2 mM hydroxyurea (HU) for 24 hours (h). pKAP1 (S824) and pCHK (S345) serve as markers of ATM and ATR signaling, respectively.

**Extended Data 6 – related to Figure 4. Further characterization of late-replicating TA-rich DSB sites: a,** A Venn diagram showing the intersection between the 72 break sites and the 125 common fragile sites in the humCFS database. **b,** Example screenshot of MAST output from the 72 break sites. The most common motifs are shown in red and light blue. Peak sequences contain multiple copies of the respective motifs. **c,** Quantification of V5-FANCM ChIP-qPCR at a TA-rich repeat and a control locus is shown. V5-FANCM expression was induced in either wild-type or *SMARCAL1* KO hTERT RPE1 *TP53* KO cells for 48 hours with 1 µg/ml doxycycline. *n* = 2 experimental replicates. **d,** Heatmap showing wavelet-smoothed signal from Repli-seq data in GM06990 cells across 72 MRE11 ChIP-Seq peaks only found in the *SMARCAL1* KO:sg*FANCM* cell. A 2 Mb region around each peak is shown and peaks are ordered ascending from late to early replication timing. **e,** Pie chart showing 72 MRE11 ChIP-Seq peaks only found in the *SMARCAL1* KO:sg*FANCM* cell categorized into replication timing phases based on the maximum signal below each peak using percentage-normalized signal tracks from Repli-seq data in GM06990 cells.

## Acknowledgements

We thank Joseph M. Replogle for helpful discussions on sgRNA selection and Bruno Silva and Claus M Azzalin for sharing pLVX-TetOne empty vector, FANCM wild-type, and K117R plasmids. We thank the Massimo Lopes laboratory for providing DNA fiber assay reagents. We thank Nadja Greter for design of the screen schematic. We thank Eric Aird, Erman Karasu, David Morales, Matthias Muhar, and other members of the Corn laboratory for assistance, discussion, and reagents. We also acknowledge the ETH Genome Engineering and Measurement Lab (GEML) for performing NGS and the Institute of Molecular Health Science Flow Cytometry Core Facility for technical support and access to FACS machines.

JF is a recipient of the EMBO Postdoctoral Fellowship (ALTF 220-2021). DNG is supported by a fellowship from the Helen Hay Whitney Foundation. MRDS is a recipient of the EMBO Postdoctoral Fellowship (ALTF 710-2021). MFS is a recipient of the Boehringer Ingelheim Fonds (BIF) PhD fellowship. The research in the Cejka laboratory is supported by ERC (101018257) and Swiss National Science Foundation grants (310030_207588 and 310030_205199). This work was supported by National Institutes of Health grant R00GM130964 to MJ. The research in the Corn laboratory was supported by the European Research Council (ERC) under the European Union’s Horizon 2020 research and innovation programme (Grant agreement No. 855741-DDREAMM-ERC-2019-SyG).

## Code availability

The modified gemini_calculate_lfc_mod function is provided as a html file in the supplementary information file.

## Data availability

The CRISPRi screen dataset is available through the NCBI BioProject database (BioProject: PRJNA988447). ChIP-Seq data is available through GEO (accession code: GSE236062).

