## Supplementary Figures 1-6 & Supplementary Tables 3-4 for "Comprehensive Interrogation of Synthetic Relationships in the Human DNA Damage Response"

### Extended Data Fig. 1 - related to Fig. 1

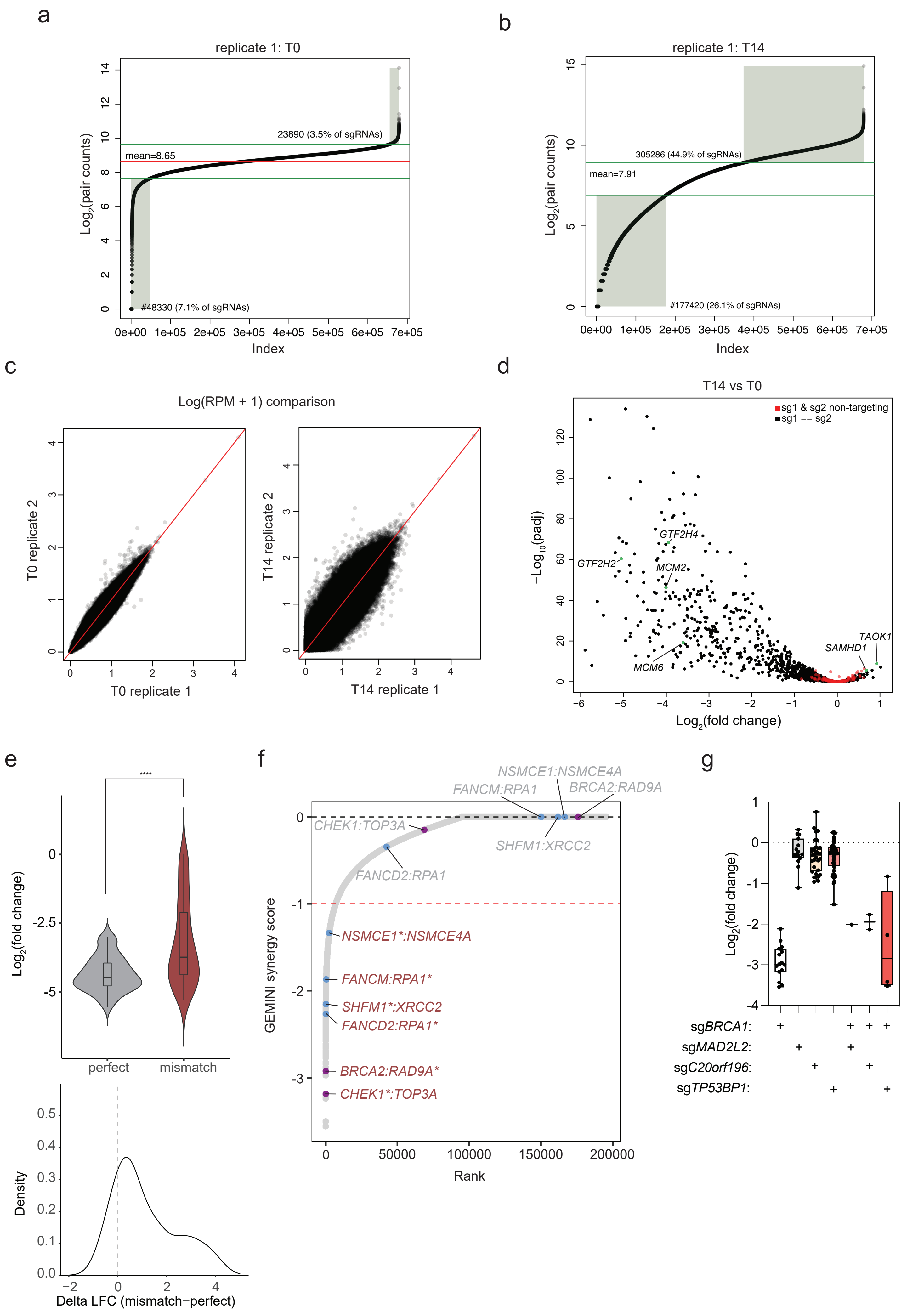

### Extended Data Fig. 2 - related to Fig. 1

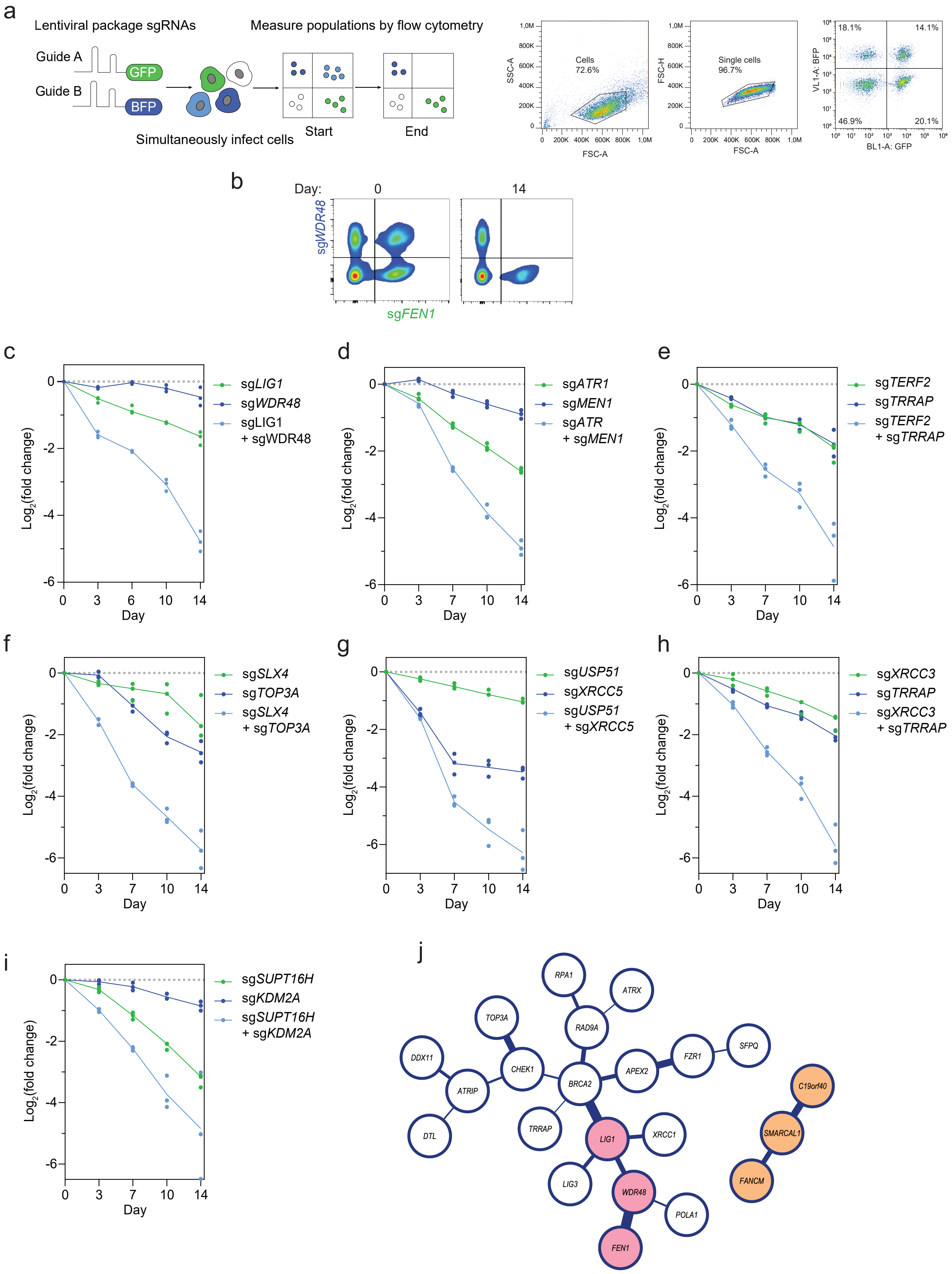

### Extended Data Fig. 3 - related to Fig. 2

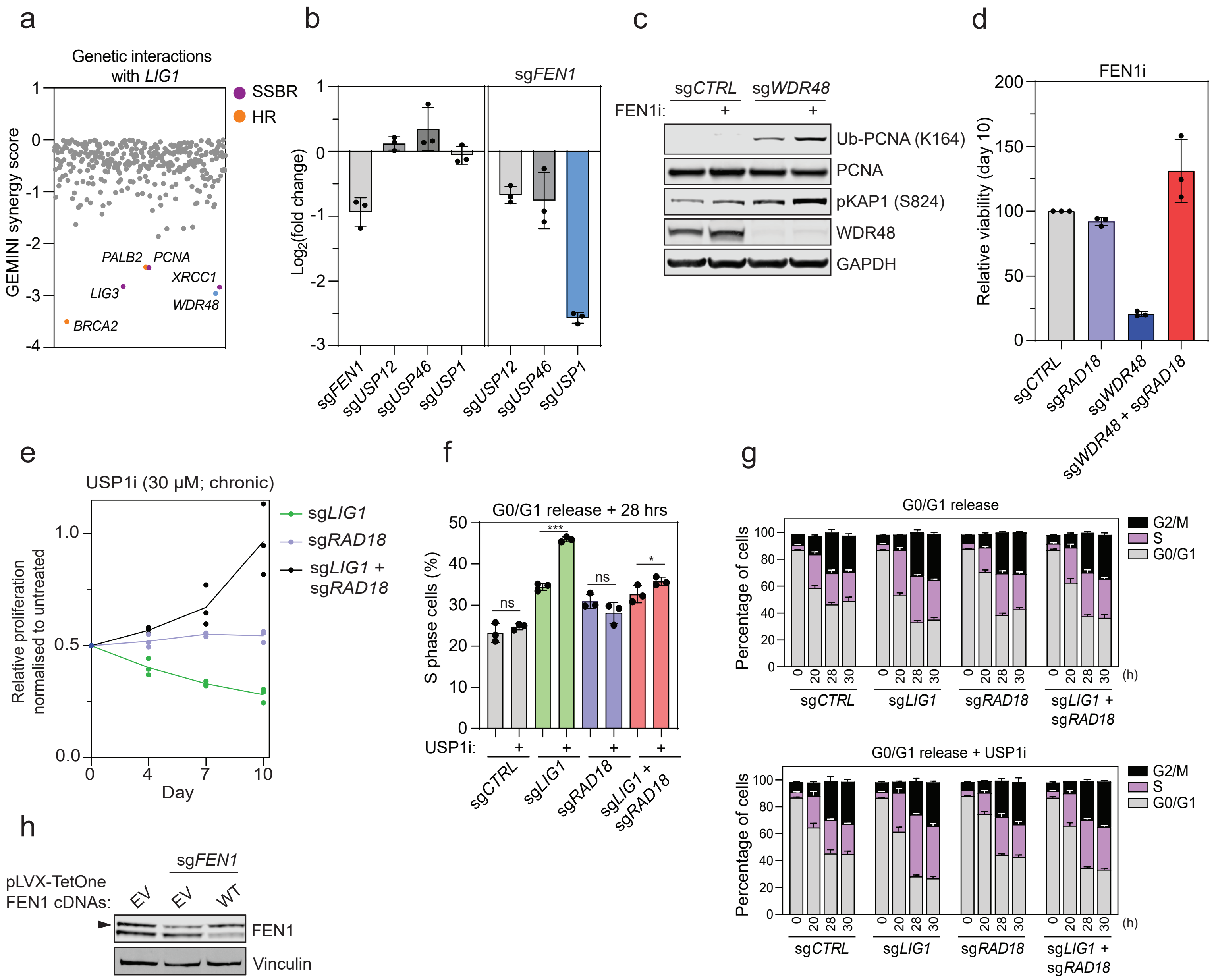

### Extended Data Fig. 4 - related to Fig. 3

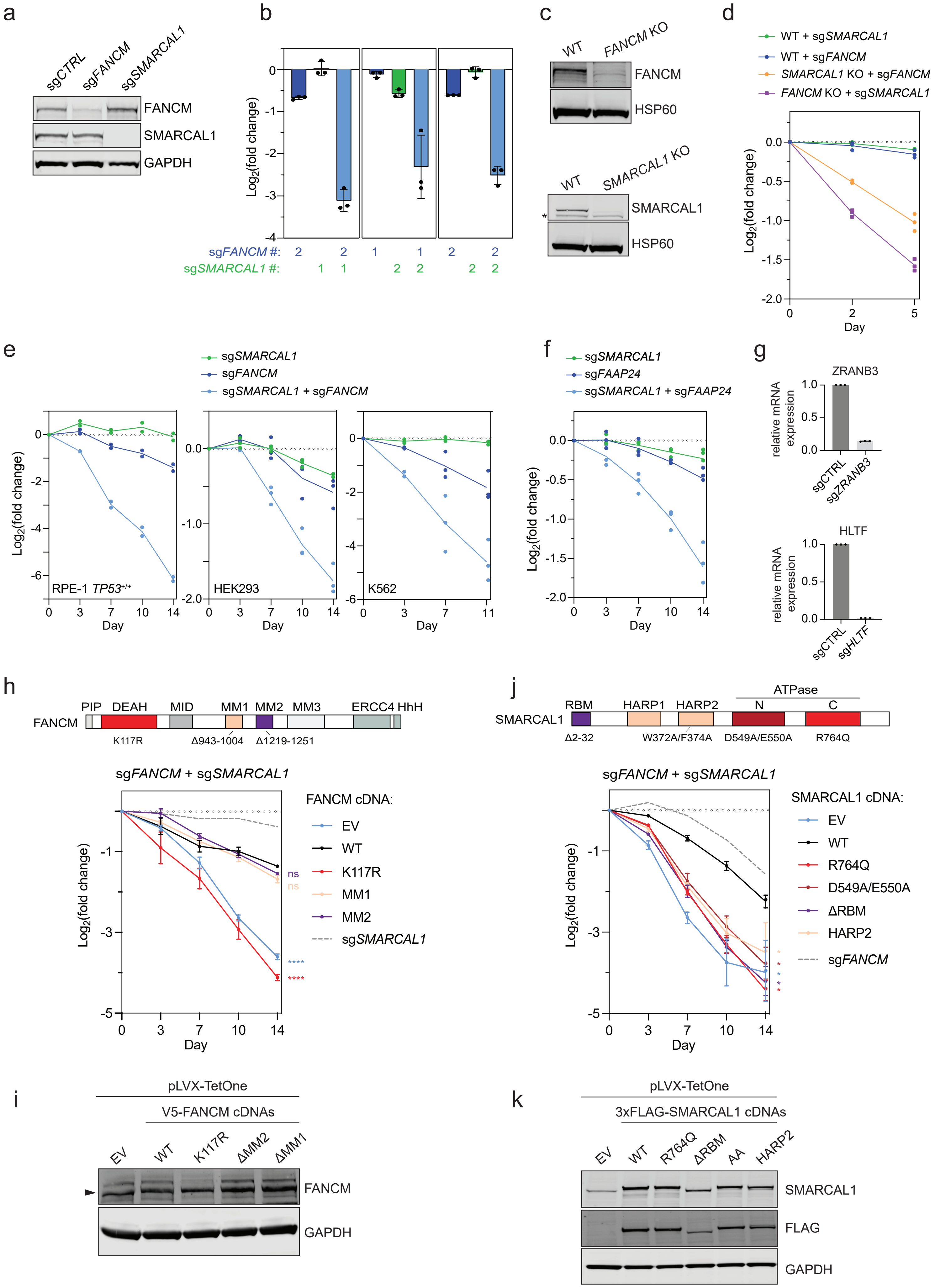

### Extended Data Fig. 5 - related to Fig. 3

a

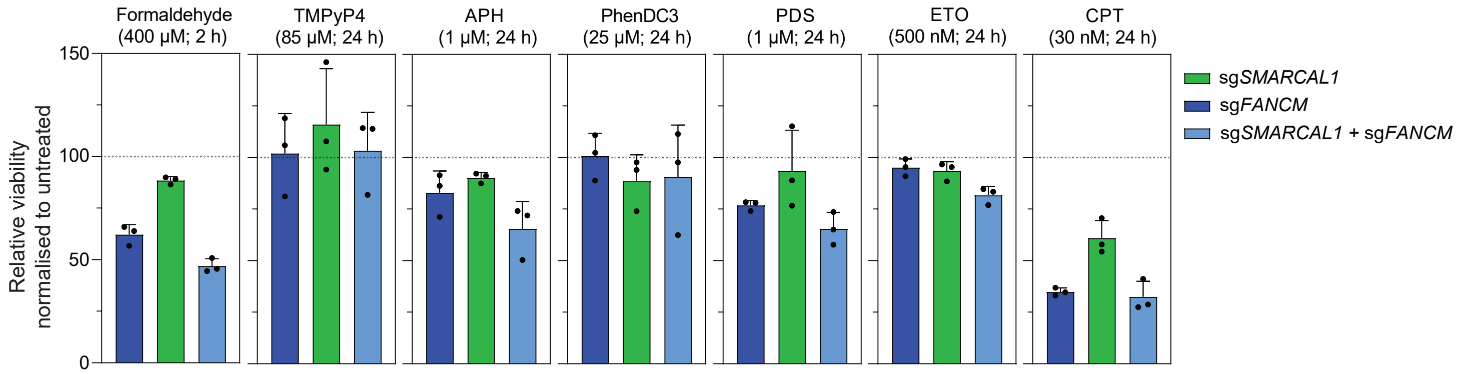


b

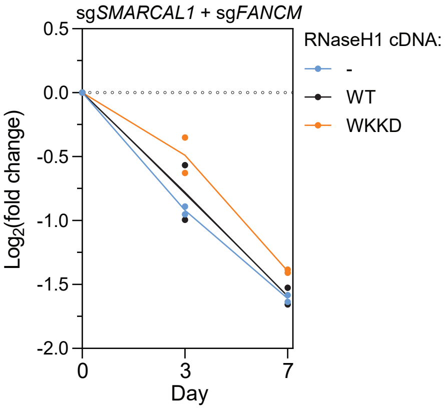


c

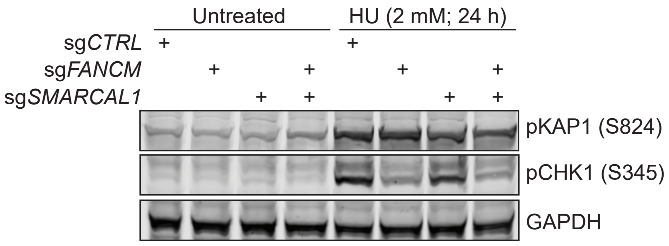

### Extended Data Fig. 6 - related to Fig. 4

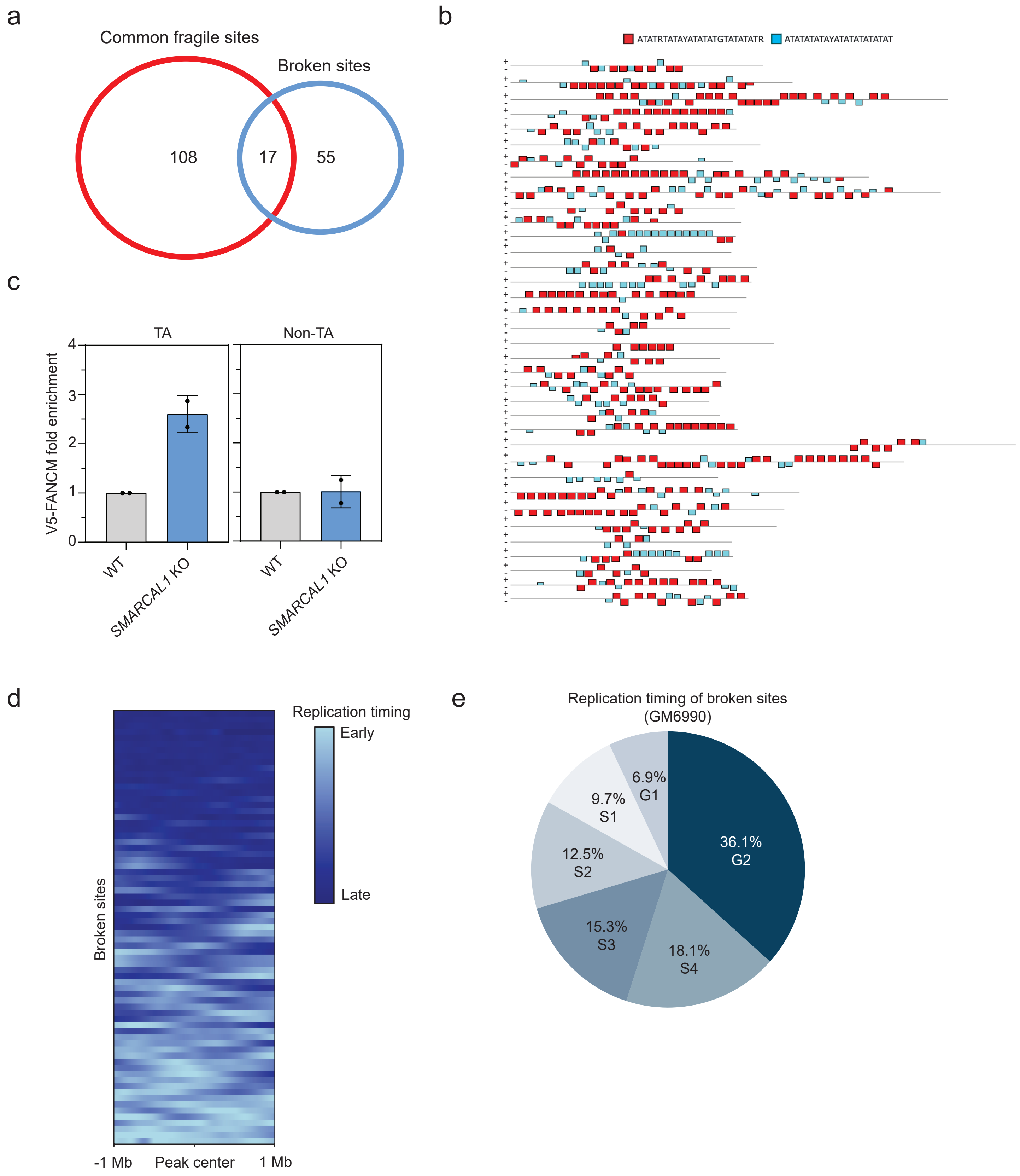

**Supplementary Table 3:** CRISPRi and CRISPR KO guide sequences used in validation studies.

| sgRNA | Protospacer sequence |
|---|---|
| FEN1 | GCGGGAGCGCGGGCTTTGGA |
| TERF2 | GATGGCCGCGGGGGCCGGGA |
| SUPT16H | GGGCGGGAAGAGACTGCGAG |
| USP51 | GGCCGATGGGAGGGGTCCTG |
| ATR | GGCCCGACGGAGCCGTGTGG |
| KDM2A | GGGTGTTGTTTCCCTCACAC |
| MEN1 | GTGCGCCGCGGTGCCTAGTG |
| C19orf40/FAAP24 | GAGGCCCGAATACAGCCGGC |
| XRCC5 | GGACCAAAGCGCCTGAGGAC |
| SMARCAL1_1 | GGTCTTCCGAGTGGACGGTT |
| SMARCAL1_2 | GGAGTGGACGGTTAGGCCAC |
| LIG1 | GCGCGAACTTGGGACTGCAG |
| SLX4 | GCCGCGGCCGCGCCGGAGGT |
| XRCC3 | GCGGCGGGCCTCACCTCCCG |
| FANCM_1 | GAGCAGCAGCTCAACCGCTA |
| FANCM_2 | GGAAGGAAACCGATGGGGAT |
| WDR48/UAF1 | GGCGGAGGAAAGTGCAGGTA |
| TRRAP | GCGGGACGGCGACCAAAGGA |
| TOP3A | GACTGGATTCGGTGCTGAGG |
| FANCD2 | GGAAAGTCGAAAACTACGGG |
| USP1 | GCCAGGAGCGCAACAAGCAG |
| USP12 | GGCCGGGAGTGCCTGAGCGC |
| USP46 | GACTGCCCATGGTGGCGCGC |
| RAD18 | GGCATCCTCGGGAGCGACCA |
| ZRANB3 | GACCTCAGGCTCCAACTCGT |
| HLTF | GTCAGGAGCGCACGACTGAA |
| FANCA | GGCCTTGGCGCCTACAGCCC |
| CTRL (non-targeting; pLG1-mCherry) | GATATCCCGGTGGGGTTCTC |
| CTRL (non-targeting; pSLQ-1371-BFP) | GTCAGTGCGGCGGGGACGAA |
| FANCM_KO | GAAATTGTACATGACCACGG |
| SMARCAL1_KO | GGAGAAGGTGAGGCCCATAT |

**Supplementary Table 4:** Key resources & reagents

| Reagent or resource | Source | Identifier |
|---|---|---|
| **Antibodies** | | |
| WDR48 | Proteintech | Cat# 16503-1-AP, RRID:AB_2878266 |
| FEN1 | Abcam | Cat# ab153825, RRID:AB_2938984 |
| KAP1 (phospho S824) | Abcam | Cat# ab70369, RRID:AB_1209417 |
| DNA Ligase I | Proteintech | Cat# 18051-1-AP, RRID:AB_2265726 |
| Rad18 (D2B8) XP® | Cell Signaling Technology | Cat# 9040, RRID:AB_2756446 |
| Ubiquityl-PCNA (Lys164) (D5C7P) | Cell Signaling Technology | Cat# 13439, RRID:AB_2798219 |
| Phospho-Histone H2A.X (Ser139) (20E3) | Cell Signaling Technology | Cat# 9718, RRID:AB_2118009 |
| Vinculin antibody [VIN-54] | Abcam | Cat# ab130007, RRID:AB_11156698 |
| 53BP1 | Novus | Cat# NB100-304SS, RRID:AB_920462 |
| FANCM | Abcam | Cat# ab95014, RRID:AB_10675719 |
| SMARCAL1 (E-12) | Santa Cruz Biotechnology | Cat# sc-376377, RRID:AB_10987841 |
| HSP60 (N-20) | Santa Cruz Biotechnology | Cat# sc-1052, RRID:AB_631683 |
| Phospho-Chk1 (Ser345) (133D3) | Cell Signaling Technology | Cat# 2348, RRID:AB_331212 |
| GAPDH, Clone D4C6R | Cell Signaling Technology | Cat# 97166, RRID:AB_2756824 |
| FLAG | Sigma-Aldrich | Cat# F3165, RRID:AB_259529 |
| MRE11 | Novus | Cat# NB100-142, RRID:AB_10077796 |
| CD55 | BioLegend | Cat# 311312, RRID:AB_2075856 |
| CldU/BrdU | Abcam | Cat# ab6326, RRID:AB_305426 |
| IdU/BrdU | BD Biosciences | Cat# 347580, RRID:AB_10015219 |
| V5 | Thermo Fisher Scientific | Cat# R96025, RRID:AB_159313 |
| Goat anti-Mouse IgG Alexa Fluor™ 488 | Thermo Fisher Scientific | Cat# A-11001, RRID:AB_2534069 |
| Goat anti-Rabbit IgG Alexa Fluor™ 488 | Thermo Fisher Scientific | # A-11008, RRID:AB_143165 |
| Goat anti-Rat IgG Alexa Fluor™ 568 | Molecular Probes | Cat# A-11077, RRID:AB_141874 |
| IRDye 800CW Donkey anti-Rabbit IgG | LI-COR Biosciences | Cat# 926-32213, RRID:AB_621848 |
| IRDye 800CW Donkey anti-Mouse IgG | LI-COR Biosciences | Cat# 926-32212, RRID:AB_621847 |
| **Chemicals, Peptides, and Recombinant Proteins** | | |
| FEN1-IN-1 | MedChem Express | Cat# HY-123834 |
| KSQ-4279 | MedChem Express | Cat# HY-145471 |
| Hydroxyurea | Sigma-Aldrich | Cat# H8627 |
| Mitomycin C | Sigma-Aldrich | Cat# M0503 |
| Etoposide | Sigma-Aldrich | Cat# E1383 |

| | | |
|---|---|---|
| Camptothecin | Sigma-Aldrich | Cat# C9911 |
| Aphidicolin | Sigma-Aldrich | Cat# A0781 |
| Formaldehyde | Thermo Fisher Scientific | Cat# 28908 |
| Olaparib | Selleck Chemicals | Cat# S1060 |
| Pyridostatin Trihydrochloride | RayBiotech | Cat# 332-11923 |
| TMPyP4 | Sigma-Aldrich | Cat# 613560 |
| JH-RE-06 | Sigma-Aldrich | Cat# SML2993 |
| ML-323 | MedChem Express | Cat# HY-17543 |
| CldU (5-chloro-2'-deoxyuridine) | MedChem Express | Cat# CS-0059203 |
| IdU (5-iodo-2′-deoxyuridine) | Astatech Inc | Cat# AST-34454 |
| EdU (5-ethynyl-2′-deoxyuridine) | Thermo Fisher Scientific | Cat# A10044 |
| G418 | Thermo Fisher Scientific | Cat# 11811031 |
| Puromycin Dihydrochloride | Thermo Fisher Scientific | Cat# A1113803 |
| Penicillin-Streptomycin | Thermo Fisher Scientific | Cat# 15-140-122 |
| Fetal Bovine Serum | Gibco | Cat# 10270106 |
| DMEM/F-12, GlutaMAX™ supplement | Thermo Fisher Scientific | Cat# 10565018 |
| cOmplete™, EDTA-free Protease Inhibitor Cocktail | MilliporeSigma | Cat# 11873580001 |
| Halt™ Phosphatase Inhibitor Cocktail | Thermo Fisher Scientific | Cat# 78426 |
| Polybrene | Sigma-Aldrich | Cat# H9268 |
| Trypsin EDTA (0.25%), Phenol red | Thermo Fisher Scientific | Cat# 25-200-056 |
| ProLong™ Gold Antifade Mountant with DAPI | Thermo Fisher Scientific | Cat# P36931 |
| ProLong™ Diamond Antifade Mountant | Thermo Fisher Scientific | Cat# P36965 |
| Colcemid | AdipoGen | Cat# AGCR13567M005 |
| Phosphate Buffer Saline (PBS) | Gibco | Cat# 10010023 |
| S1 nuclease | Thermo Fisher Scientific | Cat# EN0321 |
| DAPI solution | BD Biosciences | Cat #564907 |
| Blasticidin S | Sigma-Aldrich | Cat #SBR00022 |
| Hoechst 33342 Ready Flow™ Reagent | Thermo Fisher Scientific | Cat # R37165 |
| SsoAdvanced Universal SYBR® Green Supermix | Bio-Rad | Cat # 1725270 |
| iScript™ Reverse Transcription Supermix | Bio-Rad | Cat # 1708840 |

| | | |
|---|---|---|
| **Critical Commercial Assays** | | |
| Gentra Puregene Cell Kit | QIAGEN | Cat# 158767 |
| NEBNext Ultra II Q5 Master Mix | New England Biolabs | Cat# M0544 |
| NEB Hi-Fi Assembly Mastermix | New England Biolabs | Cat# E2621 |
| P3 Primary Cell 4D-Nucleofector® X Kit L | Lonza | Cat# V4XP-3024 |
| RNeasy Kit | QIAGEN | Cat# 74104 |
| **Experimental Models: Cell Lines** | | |
| hTERT RPE1 dCas9-KRAB TP53−/− | This study | N/A |
| hTERT RPE1 dCas9-KRAB TP53+/+ | This study | N/A |
| K562 dCas9-KRAB | This study | N/A |
| HEK293 dCas9-KRAB | This study | N/A |
| **Oligonucleotides** | | |
| Protospacer sequences for all CRISPR KO or CRISPRi experiments | See Supplementary Table 3 | N/A |
| PCR primers for screen sequencing | Replogle et al., 2022 | oJR232: AATGATACGGCGACCACCGAGATCTACACTCTTTCCCTACACGACGCTCTTCCGATCTgtatcccttggagaaCCAcctTGTTG<br>oJR233: CAAGCAGAAGACGGCATACGAGATtcgccttaGTCTCGTGGGCTCGGAGATGTGTATAAGAGACAGCTATGCTGTTTCCAGCtTAGCTCTtAAAC |
| qPCR primers | | GAPDH fwd: CAACAGCGACACCCACTCCT<br>GAPDH rev: CACCCTGTTGCTGTAGCCAAA<br>HLTF fwd: CGTATTAGAGAACCGGCCTTAC<br>HLTF rev: GGATCACTCTTAGCCACCTTATG<br>ZRANB3 fwd: GGTGTATGGTGGCTGATGAA<br>ZRANB3 rev: AGAGACGAAGGGACCACTATTA<br>ChIP_TA fwd: TTGTGGCAAACAAAATACAA<br>ChIP_TA rev: GCCAAACCATATCACTAACT<br>ChIP_non-TA fwd: GGAAACTTCTAGTGCTTCAA<br>ChIP_non-TA rev: TTGTCATACGTCTGAGTCAG |

| | | |
|---|---|---|
| **Recombinant DNA** | | |
| pLVX-TetOne-Puro | Silva et al., 2019 | N/A |
| pLVX-TetOne-Puro-V5-FANCM_WT | Silva et al., 2019 | N/A |
| pLVX-TetOne-Puro-V5-FANCM_K117R | Silva et al., 2019 | N/A |
| pLVX-TetOne-Puro-V5-FANCM_ΔMM1 | This study | N/A |
| pLVX-TetOne-Puro-V5-FANCM_ΔMM2 | This study | N/A |
| pLVX-TetOne-Puro-3xFLAG-SMARCAL1_WT | This study | N/A |
| pLVX-TetOne-Puro-3xFLAG-SMARCAL1_R764Q | This study | N/A |
| pLVX-TetOne-Puro-3xFLAG-SMARCAL1_ΔRBM | This study | N/A |
| pLVX-TetOne-Puro-3xFLAG-SMARCAL1_ΔHARP | This study | N/A |
| ppyCAG_RNaseH1_WT | Xiang-Dong Fu lab | Addgene plasmid #111906 |
| ppyCAG_RNaseH1_WKKD | Xiang-Dong Fu lab | Addgene plasmid #111905 |
| ppyCAG_RNaseH1_WT_IRES_mCherry | This study | N/A |
| ppyCAG_RNaseH1_WKKD_IRES_mCherry | This study | N/A |
| pShuttle-FEN1hWT | Sheila Stewart lab | Addgene plasmid #35027 |
| pLVX-TetOne-Puro-FEN1_WT | This study | N/A |
| **Software and Algorithms** | | |
| LI-COR ImageStudio software V5.2 | LI-COR | https://www.licor.com/bio/image-studio/ |
| Adobe Illustrator 2023 | Adobe Systems | https://creativecloud.adobe.com/en/apps/download/creative-cloud |
| Prism 9 | GraphPad | https://www.graphpad.com/scientific-software/prism/ |
| ImageJ | NIH | https://imagej.net/ij/index.html |
| FlowJo_v10.8.1 | FlowJo | https://www.flowjo.com/solutions/flowjo/downloads |
| GEMINI | Zamanighomi et al., 2019 | https://github.com/sellerslab/gemini |
| Cytoscape 3.9.1 | Cytoscape | https://cytoscape.org/ |
| ICE CRISPR analysis tool | Synthego | https://www.synthego.com/products/bioinformatics/crispr-analysis |
| R 4.2.1/2 | R Core Team | https://www.R-project.org/ |
